# FHOD-1 is the only formin in *Caenorhabditis elegans* that promotes striated muscle growth and Z-line organization in a cell autonomous manner

**DOI:** 10.1101/2020.06.18.159582

**Authors:** Sumana Sundaramurthy, SarahBeth Votra, Arianna Laszlo, Tim Davies, David Pruyne

## Abstract

The striated body wall muscles of *Caenorhabditis elegans* are a simple model system with well-characterized sarcomeres that have many vertebrate protein homologs. Previously, we observed deletion mutants for two formin genes, *fhod-1* and *cyk-1*, developed thin muscles with abnormal dense bodies/sarcomere Z-lines. However, the nature of the *cyk-1* mutation necessitated maternal CYK-1 expression for viability of the examined animals. Here, we tested the effects of complete loss of CYK-1 using a fast acting temperature-sensitive *cyk-1(ts)* mutant. Surprisingly, neither post-embryonic loss of CYK-1 nor acute loss of CYK-1 during embryonic sarcomerogenesis caused muscle defects, suggesting CYK-1 might not play a direct role in muscle development. Consistent with this, examination of *cyk-1(*Δ) mutants re-expressing CYK-1 in a mosaic pattern showed CYK-1 cannot rescue muscle defects in a muscle cell autonomous manner, suggesting muscle phenotypes caused by *cyk-1* deletion are likely indirect. Conversely, mosaic re-expression of FHOD-1 in *fhod-1(Δ)* mutants promoted muscle cell growth, as well as proper Z-line organization, in a muscle cell autonomous manner. As we can observe no effect of loss of any other worm formin on muscle development, we conclude that FHOD-1 is the only formin that directly promotes striated muscle development in *C. elegans*.

## Introduction

Striated muscles, found widely across the animal kingdom (Clark, McElhinny, Beckerle, & Gregorio, 2002), are so-called due to the appearance of striations composed of regularly repeating contractile units, called sarcomeres. Each sarcomere is bordered by Z-lines that anchor actin-based thin filaments, while myosin-based thick filaments interdigitate between thin filaments and are anchored at the M-line at the sarcomere center. Besides these major cytoskeletal components, the contractile lattice is made up of various other sarcomeric proteins that help maintain the structure and function of muscle (Henderson, Gomez, Novak, Mi-Mi, & Gregorio, 2017).

Evidence from multiple model systems demonstrates that formins contribute to the assembly of sarcomeres. Unique among stimulators of actin assembly, many formins both nucleate and promote elongation of actin filaments in the presence of profilin-bound G-actin, while remaining associated with the barbed end of the growing actin filament (Pruyne et al., 2002; Sagot, Rodal, Moseley, Goode, & Pellman, 2002; Pring, Evangelista, Boone, Yang, & Zigmond, 2003; Moseley et al., 2004; Kovar & Pollard, 2004; Romero et al., 2004; Kovar, Harris, Mahaffy, Higgs, & Pollard, 2006). Formins have two conserved Formin Homology (FH) domains, the FH1 and FH2. When dimerized, FH2 domains nucleate and protect the growing barbed-end from inhibitors of elongation, like capping proteins (Pruyne et al., 2002; Sagot et al., 2002; Pring et al., 2003; Kovar & Pollard, 2004; Moseley et al., 2004). The proline-rich FH1 domain recruits profilin bound to G-actin, which allows the addition of G-actin to the barbed ends (Chang, Drubin, & Nurse, 1997; Evangelista et al., 1997; Imamura et al., 1997; Romero et al., 2004; Kovar et al., 2006).

Based on sequence homology, animal formins are grouped into nine families (Higgs & Peterson, 2005; Pruyne, 2016). One member of the FHOD-family of formins, FHOD3, is the best-studied vertebrate formin for its effects on muscle development. Variants of the *fhod3* gene have been associated with occurrence of hypertrophic and dilated cardiomyopathies in humans (Arimura et al., 2013; Wooten et al., 2013; Hayashi et al., 2018; Ochoa et al., 2018), while knock out of *fhod3* in mice results in lethality at embryonic day 11.5 due to cardiac insufficiency, with malformed sarcomere-containing myofibrils with immature Z-lines in the cardiac muscle (Kan-O et al., 2012). This crucial role of FHOD3 depends on its ability to interact with actin (Fujimoto et al., 2016). A conditional knock out in mice showed postnatal loss of FHOD3 causes only mild impairment to cardiac function, and induced upregulation of fetal cardiac genes, suggesting that FHOD3 is not essential for sarcomere maintenance, but might modulate hypertrophic changes (Ushijima et al., 2018). Studies with human induced pluripotent stem cell-derived cardiomyocytes and rat cardiomyocytes confirm FHOD3 regulates myofibrillogenesis (Taniguchi et al., 2009; Iskratsch et al., 2010; Fenix et al., 2018). Studies in *Drosophila* indirect flight muscles (IFMs) have shown lack of the fly’s only FHOD-family formin FHOS abolished sarcomere organization, while early knockdown led to thin, irregular myofibrils with dispersed and rudimentary-appearing Z-discs, and late knockdown prevented thin filament elongation or new thin filament incorporation as sarcomeres enlarged (Shwartz, Dhanyasi, Schejter, & Shilo, 2016).

Formins of the DAAM-family have also been implicated in muscle development. In *Drosophila* IFMs, absence of DAAM led to thin myofibrils with reduced F-actin and thick filament content, and partially disorganized sarcomeres with abnormal Z-discs, M-lines, and thick and thin filaments (Molnar et al., 2014). Similar, but lesser, defects were also observed in heart and larval body wall muscles partially deficient for DAAM (Molnar et al., 2014). In mice, conditional DAAM1 knock out led to non-compaction cardiomyopathy with misshapen hearts and poor cardiac function, and simultaneous knock out of DAAM1 and DAAM2 led to even stronger myopathy, with severely reduced cardiac function and disrupted sarcomere structure (Ajima et al., 2015). An RNA interference (RNAi)-based study in neonatal mouse cardiomyocytes confirmed the importance of DAAM1 for myofibril organization, and also implicated FMNL-family formins FMNL1 and FMNL2, as well as the DIAPH-family formin DIAPH3 (Rosado et al., 2014). While these studies suggest formins promote sarcomere assembly in striated muscles, it is not clear what roles they play.

Our work with the simple model nematode *Caenorhabditis elegans* has implicated formins in promoting sarcomere formation in its striated muscle as well. The worm has two pairs of body wall muscles (BWMs) that extend the length of the animal in dorsal and ventral positions. Each BWM cell consists of an organelle-containing cell body overlying a spindle-shaped myofilament lattice that is adherent to a basal lamina. Distinct from cross-striated vertebrate or fly muscle, BWM cells are obliquely striated due to the orientation of striations at a 6° angle with respect to the longitudinal axis of the cell. The majority of thin filaments are anchored at dense bodies (DBs), which are analogous to vertebrate Z-lines, whereas thick filaments are attached to the M-line (Moerman & Williams, 2006; Gieseler et al., 2017). In BWM, all DBs and M-lines are also attached to the plasma membrane, and based on protein composition, the DBs also resemble costameres and focal adhesions (Lecroisey, Ségalat, & Gieseler, 2007).

The sarcomeric protein composition and interactions in BWM are similar to mammalian striated muscle, with many vertebrate homologs (Benian & Epstein, 2011). But unlike mammals, where fifteen genes represent seven formin families, *C. elegans* has only six formin genes (*fhod-1, cyk-1, daam-1, frl-1, exc-6* and *inft-2*), which represent five families (Pruyne, 2016). Due to this simplicity of *C. elegans*, its BWM serves as a very good model to study roles of formins in the development and function of striated muscle.

We previously characterized deletion mutants for five of the worm formin genes for defects in striated BWMs (Mi-Mi et al., 2012; Mi-Mi & Pruyne, 2015). Among these, we observed loss of only the FHOD-family FHOD-1 and the DIAPH-family CYK-1 resulted in muscle defects (Mi-Mi et al., 2012). Based on the tendency of formins to promote assembly of long, unbranched actin filaments *in vitro*, an attractive model was that FHOD-1 and CYK-1 initiate the assembly of the actin-based thin filaments, with the prediction that complete absence of both formins should prevent thin filament formation. Our analysis was based on strains with deletion alleles that eliminated part or all of the actin-interacting FH2 domains coded in *fhod-1* and *cyk-1*, making them putative nulls for formin-mediated actin assembly. Loss of either formin individually resulted in thin BWM cells with fewer striations per cell (Mi-Mi et al., 2012). The *fhod-1(Δ); cyk-1(Δ)* double mutants had even thinner BWMs, but those still had sarcomeres with thin filaments, inconsistent with our preliminary prediction that thin filaments would be absent (Mi-Mi et al., 2012; Mi-Mi & Pruyne, 2015). However, CYK-1 is also essential for cytokinesis during embryonic development (Swan et al., 1998), and thus homozygous *cyk-1* mutants had to be derived from heterozygous parents, leaving the possibility that maternally inherited CYK-1 supported sarcomere assembly in the double mutant. The recent development of a fast acting temperature-sensitive allele of *cyk-1* provides us an opportunity to re-examine the contribution of this formin to muscle development (Davies et al., 2014).

## Results

### Post-embryonic CYK-1 loss leads to minimal BWM developmental defects

To characterize the effects of loss of FHOD-1 and CYK-1, we had previously used RNAi as well as strains with deletion alleles *fhod-1(tm2363)* and *cyk-1(ok2300)*, referred to hereafter as *fhod-1(Δ)* and *cyk-1(Δ)*, respectively (Mi-Mi et al., 2012; Mi-Mi & Pruyne, 2015). However, *cyk-1(*Δ) requires use of mutants that are maternally-rescued, and *cyk-1(RNAi)* has a relatively slow onset. In contrast, the novel fast acting temperature-sensitive allele *cyk-1(or596ts)* produces a CYK-1 product that supports embryonic development at a permissive temperature of 16°C, but becomes inactive within 5 min after shift to 26°C (Davies et al., 2014). We used this *cyk-1(ts)* in combination with *fhod-1(Δ)* to analyze the effects of complete loss of CYK-1 and FHOD-1 on post-embryonic BWM development.

Age synchronized L1 stage larvae were allowed to grow at either permissive or restrictive temperature, and samples were collected every 12 hours and stained with fluorescently labeled phalloidin to visualize filamentous actin (F-actin) to track BWM growth. Even prior to temperature shift, approximately 10% *cyk-1(ts)* and *fhod-1(Δ)* single mutants and 20% *fhod-1(Δ); cyk-1(ts)* double mutants had an apparent failure in body elongation during embryogenesis, an effect related to formin functions in the epidermis, as described previously for *fhod-1* and *cyk-1* (Vanneste, Pruyne, & Mains, 2013; Refai, Smit, Votra, Pruyne, & Mains, 2018). As these worms died soon after plating, they were excluded from analysis. For remaining animals, total body width (as a measure of overall body size), BWM width, and individual BWM cell widths (based on F-actin stain) were measured (Fig. 1, Fig. S1).

**Figure 1.**
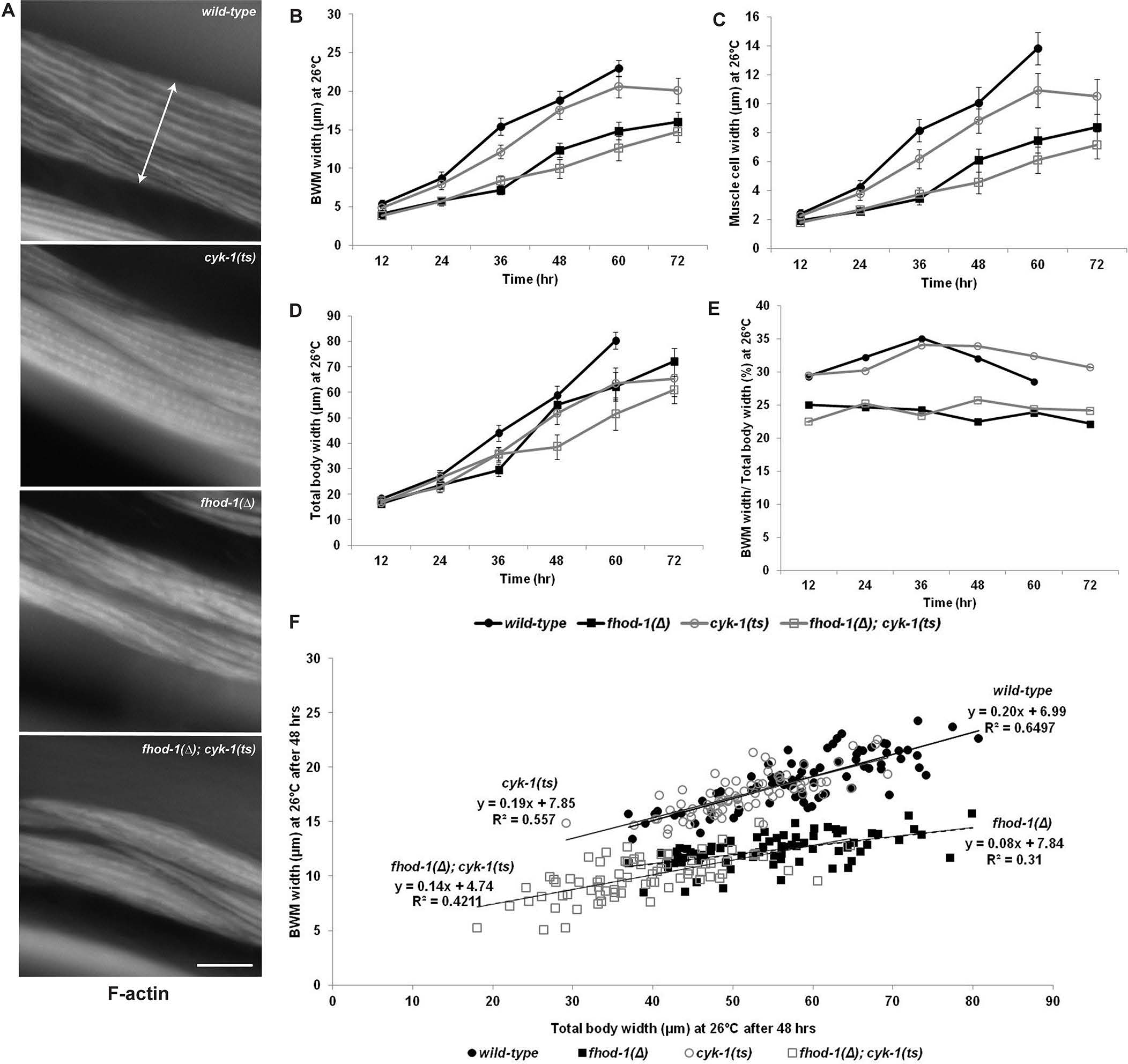
Post-embryonic CYK-1 loss leads to minimal BWM developmental defects. Larvae were hatched at permissive temperature (16°C), and grown at restrictive temperature (26°C) for up to 72 hrs, with samples collected every 12 hrs and stained with fluorescently-labeled phalloidin. (A) Dorsal views of portions of BWM in worms of indicated genotypes, grown at restrictive temperature 60 hrs. Double arrow shows lateral width of one BWM. Scale bar, 20 µm. (B) BWM widths, (C) individual muscle cell widths, and (D) total body widths were measured. Graphs depict average of the means of three experiments (n = 25 animals per strain in each experiment, with width measurement of one body, two BWMs, and four muscle cells per animal). Error bars indicate the standandard error of the mean. Differences for all pairwise comparisions at a given time point were statistically significant (p < 0.001), except for difference in total body width between *fhod-1(Δ); cyk-1(ts)* and *cyk-1(ts)* at 36 hrs, which was not significant (p > 0.05). (E) Calculated ratio of (B) average BWM width to (D) average total body width. (F) Plot of BWM width versus total body width for 75 individual worms of each genotype, grown at restrictive temperature 48 hrs, with linear trendlines shown. Wild type and *cyk-1(ts)* animals show nearly identical linear relationships between total body and BWM width, suggesting the smaller BWM size of *cyk-1(ts)* animals may be a secondary consequence of a smaller body size. Approximately linear relationships between BWM size and total body size seen for *fhod-1(*Δ) and *fhod-1(*Δ*); cyk-1(ts)* animals were similar to each other, but with consistently smaller muscle-to-body ratios than wild-type.

At permissive temperature, *cyk-1(ts)* mutants grew normally, whereas at restrictive temperature, they laid no eggs and often developed protruding vulvas, phenotypes we previously observed in *cyk-1(Δ)* mutants (Mi-Mi et al., 2012). As expected, the BWM width of wild type, *fhod-1(Δ), cyk-1(ts)*, and *fhod-1(Δ); cyk-1(ts)* mutant animals increased over time at permissive temperature as the animals grew. Also as expected, BWMs grew more slowly in *fhod-1(*Δ) animals due to slower growth of individual muscle cells (Fig. S1 A, B). In *cyk-1(ts)* mutants, we also observed narrower BWMs and narrower individual muscle cells compared to wild-type at permissive temperatures, although this effect was not as great as for *fhod-1(*Δ) animals (Fig. S1 A, B), suggesting the *cyk-1(ts)* allele may be partially non-functional at 16°C. At the restrictive temperature, BWM and individual muscle cells of *fhod-1(Δ*) mutants again grew slower than wild-type, as expected (Fig. 1 A-C). Surprisingly, BWM and muscle cells of *cyk-1(ts)* were only minimally reduced for growth compared to wild-type, and those of *fhod-1(Δ*); *cyk-1(ts)* double mutants were similar to *fhod-1(Δ*) mutants (Fig. 1 A-C), an effect much weaker than we had observed for *cyk-1(*Δ) (Mi-Mi et al., 2012).

We considered whether loss of formin might affect overall body growth, and thus effects on BWM growth might be indirect. Indeed, absence of FHOD-1 very modestly reduces overall body growth, and *cyk-1(*Δ) animals are significantly smaller than wild-type (Mi-Mi et al., 2012; Refai et al., 2018). Thus, we measured the overall body widths of worms (Fig.1 D, S1 C), and determined the ratios of the widths of BWMs to the total body. This average ratio was fairly constant for wild-type worms throughout growth, and was similar between *cyk-1(ts)* and wild-type (Fig. 1 E). Interestingly ratios for *fhod-1(Δ*) and *fhod-1(*Δ*); cyk-1(ts)* double mutants were also similar to each other, but proportionately smaller than wild-type at all ages (Fig. 1 E). To determine whether this relationship held for individual worms, as opposed to average results for a population, we plotted BWM and total body widths for individual animals and observed nearly identical linear relationships between muscle and body size for wild type and *cyk-1(ts)* animals, while those for *fhod-1(Δ)* and double mutants were different from wild-type, but similar to each other (Fig. 1 F). These data confirm absence of FHOD-1 disrupts BWM growth to a greater extent than overall growth. They also suggest there may be no specific effect on BWM growth after post-embryonic CYK-1 loss, but that BWM growth defects might be an indirect consequence of reduced overall growth of *cyk-1(ts)* animals.

Our previous work suggested three of the remaining four formins in *C. elegans* (EXC-6/INFT-1, INFT-2, and FRL-1) make no obvious contribution to BWM development (Mi-Mi et al., 2012). We had also observed an FH2-targeting deletion of the formin gene *daam-1* caused no obvious synthetic defects in combination with *fhod-1(*Δ) (Mi-Mi et al., 2012), but careful analysis was not performed due to a very closely linked genomic deletion that eliminated 13 genes and caused additional growth defects (Mangio, Votra, & Pruyne, 2015). In order to examine whether DAAM-1 contributes to muscle development in *C. elegans*, we used CRISPR/Cas9-based genome editing to introduce a premature stop codon followed by a frame-shift into the FH2-coding region of *daam-1*, generating the putative FH2-null allele *daam-1(ups39)*. On inspecting BWM in age-synchronized adult *daam-1(ups39)* mutants and *fhod-1(Δ*); *daam-1(ups39)* double mutants, we observed no defects in absence of DAAM-1 when compared to wild type or *fhod-1(*Δ) animals, respectively (Fig. S2). Our current data suggest that FHOD-1 might be the only formin that supports BWM growth during post-embryonic development in *C. elegans*.

### Constitutive absence of FHOD-1 or CYK-1, but not post-embryonic loss of CYK-1, disrupts dense body (DB) organization in adult BWM

DBs that serve as sarcomere Z-lines in BWM are structurally and compositionally similar to vertebrate focal adhesions and costameres, with homologs to proteins such as integrins, paxillin, talin, vinculin and α-actinin (Lecroisey et al., 2007). The most prominent feature of these complex structures is an α-actinin (ATN-1)-rich portion that extends into the contractile lattice to anchor thin filaments (Gieseler et al., 2017). By transmission electron microscopy (TEM), DBs of wild type BWM appear as distinct electron-dense finger-like protrusions, while in *fhod-1(Δ)* mutants they appear as multiple thinner electron-dense strands (Mi-Mi & Pruyne, 2015). Correlating with this, immunostain for ATN-1 reveals DBs as discrete puncta in wild type BWM, but partially dispersed in *fhod-1(*Δ) mutants (Fig. 2).

**Figure 2.**
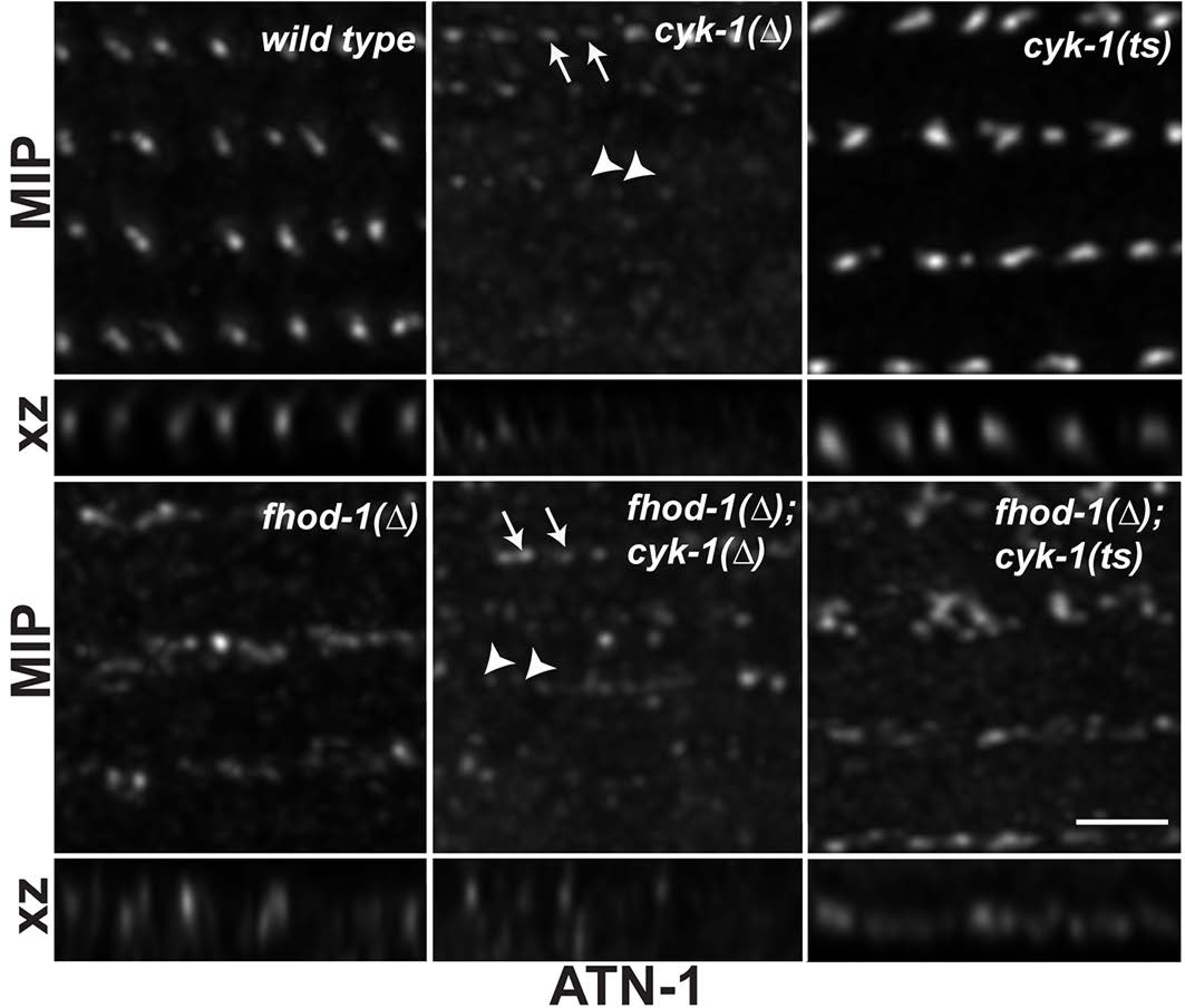
Constitutive absence of FHOD-1 or CYK-1, but not post-embryonic loss of CYK-1, disrupts DB organization. Maximum intensity projections (MIP) of dorsal views and reconstructed cross sections (xz projection) of DBs in age-synchronized adult worms after ATN-1 immunostain. Scale bar, 2 µm. DBs appear as discrete puncta in wild type BWM and appear partially dispersed in *fhod-1(*Δ) mutants. DBs of *cyk-1(Δ)* and *cyk-1(Δ); fhod-1(Δ)* mutants possess variable patterns of ATN-1 staining. Arrows point to DBs with stronger stain, and arrowheads point to regions with extremely faint stain but where we would otherwise expect DBs to be positioned. DBs of *cyk-1(ts)* appear as strongly stained puncta, and those of *fhod-1(*Δ*); cyk-1(ts)* mutants as partially dispersed by strongly stained.

By TEM, *cyk-1(Δ)* and *fhod-1(*Δ*); cyk-1(*Δ) mutant BWMs have less to almost no electron-dense material at some locations where DBs are otherwise expected (Mi-Mi & Pruyne, 2015). Consistent with this, ATN-1 immunostain of adult *cyk-1(Δ)* mutants and *fhod-1(*Δ*); cyk-1(*Δ) double mutants revealed a variable staining pattern (Fig. 2). There were some BWM regions with prominently stained structures (arrows), either wild-type-appearing puncta in *cyk-1(*Δ) mutants, or *fhod-1(*Δ)-like dispersed structures in *cyk-1(Δ); fhod-1(Δ)* double mutants. However, in other regions there were no observable ATN-1-containing structures (arrowheads), reministent of absence of DB material observed by TEM. Since regions of reduced staining were only observed in worms with *cyk-1(Δ)* in their genetic background, and we observed diffuse background stain in these regions, we reasoned this did not reflect failure of antibody penetration.

Because our results with *cyk-1(ts)* suggested postembryonic loss of CYK-1 does not affect BWM cell growth, we examined whether post-embryonic loss of CYK-1 had a similar lack of effect on DB morphology. Thus, L1 larvae were shifted to the restrictive temperature and grown 72 hrs before immunostaining for ATN-1. In contrast to *cyk-1(*Δ) mutants, DBs in *cyk-1(ts)* mutants were consistently similar to those in wild type animals grown under the same conditions, with no areas lacking ATN-1 stain (Fig. 2). Similarly, DBs of *fhod-1(Δ); cyk-1(ts)* mutants consistently appeared partially dispersed, as in *fhod-1(Δ)* mutants, with no deficient areas seen (Fig. 2). Our data suggest that constitutive absence of either FHOD-1 or CYK-1, but not post-embryonic loss of CYK-1, results in abnormal DB morphology.

### Absence of FHOD-1 but not acute loss of CYK-1 partially disrupts sarcomere structure in embryonic BWM

Lack of major defects in BWM due to post-embryonic loss of CYK-1 led us to hypothesize that the more severe defects observed in *cyk-1(Δ)* mutants were due to CYK-1 absence during embryonic BWM development. To test this hypothesis, we used the *cyk-1(ts)* strain to induce acute CYK-1 loss during embryonic myogenesis. To allow the visualization of BWMs in all embryonic stages, we crossed into the formin mutant backgrounds a transgene expressing GFP-tagged myosin heavy chain A (GFP::MYO-3) (Campagnola et al., 2002), which is expressed in BWM.

The development of embryonic BWMs is reflected in characteristic changes in appearance of muscle components in particular embryonic developmental stages (Fig. 3) (Hresko et al., 1994). After gastrulation, *C. elegans* embryos develop through stages called pre-bean, bean, comma, 1.5-fold, 2-fold, and 3-fold, named after the changing shape of the embryo. At 290 minutes after first cell division, equivalent to a pre-bean stage, myoblasts arise and are localized laterally, and gradually accumulate diffuse myosin. By 350 minutes, the comma stage, myoblasts complete migration to the dorsal and ventral quadrants, and muscle components start to polarize at the cell edges towards the hypodermis (Fig. 3, Fig. 4 A). At 420 minutes, the 1.5-fold stage, muscle cells flatten and F-actin and myosin assemble into the contractile lattice, with a polarized accumulation at cell-cell junctions between myoblasts (Fig. 3). At about 450 minutes, the 2-fold stage, we observe the first functional sarcomeres (Fig. 3), and muscle-muscle and muscle-hypodermis junctions assemble. In stages older than 2-fold, neatly aligned striations have assembled in the embryonic BWM (Fig. 3, Fig. 4 A).

**Figure 3.**
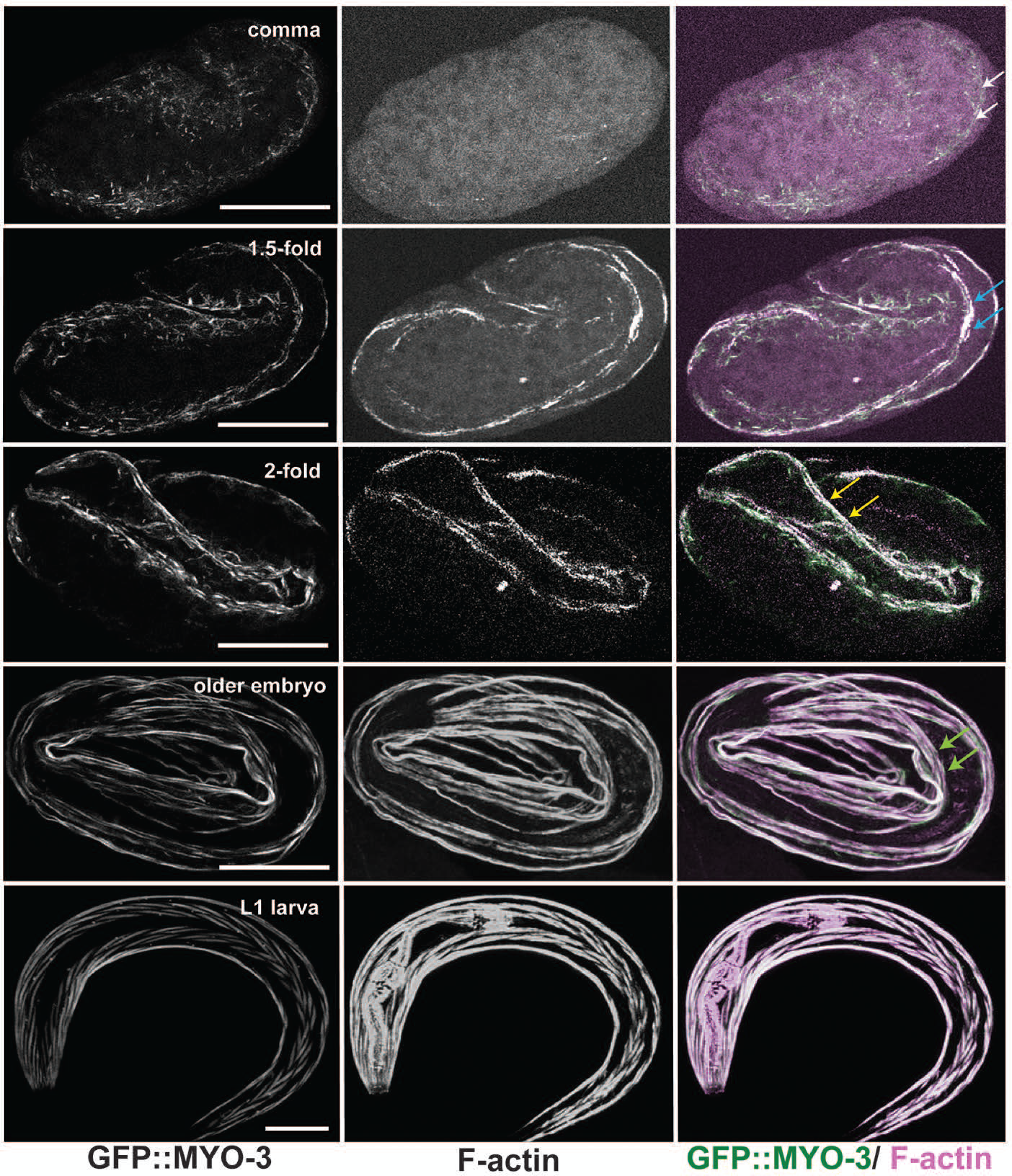
Development of the myofilament lattice in wild type embryonic BWM. Maximum intensity projections (MIP) of embryos expressing GFP::MYO-3 (a BWM marker) and stained with fluorescently-labeled phalloidin to visualize F-actin, show BWM development at embryonic stages comma, 1.5-fold, 2-fold, older than 2-fold, and L1 larva. At comma stage, traces of F-actin appear as polarization of muscle components begins (white arrows). Polarization continues through the 1.5-fold stage, where F-actin and GFP::MYO-3 strongly accumulate (blue arrows). The first sarcomeres form just prior to the 2-fold stage with appearance of defined striations (yellow arrows). At stages older than 2-fold (older embryo, L1 larva), we observe neatly aligned striations (green arrows) in BWM. Scale bars, 20 µm.

**Figure 4.**
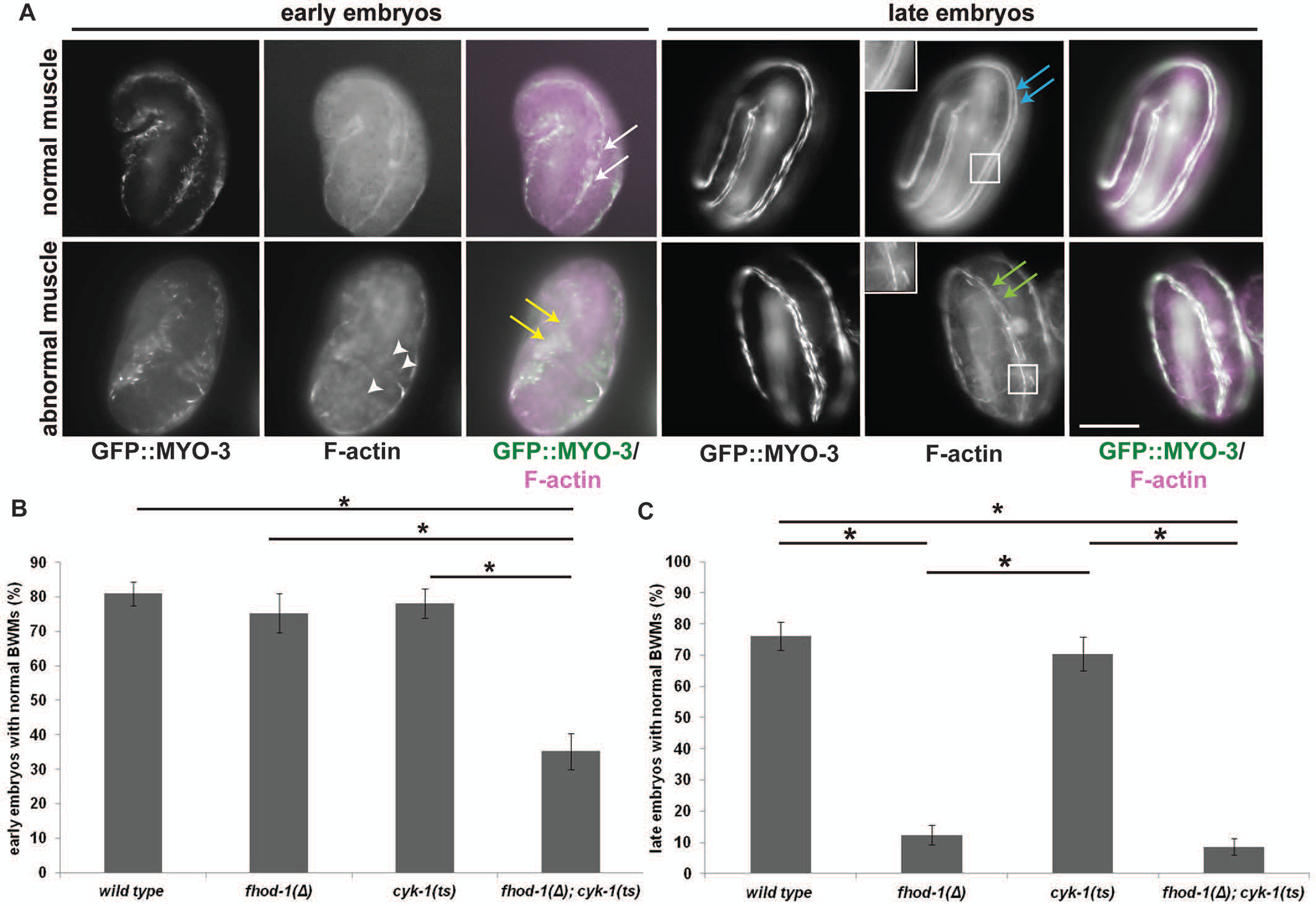
Absence of FHOD-1 but not acute loss of CYK-1 causes frayed F-actin striations in late embryonic BWM. Embryos expressing GFP-myosin grown at permissive temperature were shifted to restrictive temperature for 30 min before staining with fluorescently-labeled phalloidin to observe F-actin. (A) Normally, in early embryos (comma to 1.5-fold stage) polarization of sarcomere components is observable (white arrows), and in later embryos (2-fold and older stages) sarcomere components appear in neat striations (blue arrows). A subset of embryos of all genotypes appeared abnormal, with early embryonic stage sarcomere components failing to polarize (yellow arrows), or later embryonic stage striations appearing frayed (green arrows). Additionally, F-actin-rich nuclei (further characterized in Fig. S4) were apparent in animals bearing *cyk-1(ts)*, with or without *fhod-1(*Δ). Scale bar, 20 µm. Insets show higher magnification of late embryo striations. Quantification of early (B) and late (C) embryo phenotypes shows early defects predominate in *fhod-1(*Δ*); cyk-1(ts)* double mutants, while the later defect of fayed striations correlates with *fhod-1(*Δ), irrespective of *cyk-1*. Data shown are the averages of the means of three experiments (n = 35 early or 70 late embryos, per strain per experiment). Error bars indicate standandard error of the mean. (*) indicates p < 0.01. Differences in all other pairwise comparisons were not statistically significant (p > 0.05).

To examine embryonic myoblasts in the formin mutants, we obtained pools of mixed-stage embryos by bleaching adult worms, which liberates embryos from the adult bodies. We allowed this asynchronous population to continue developing at permissive temperature until the percentage of embryos at the two-fold stage peaked in each individual pool. Embryos were then maintained a further 30 min either at permissive or restrictive temperature before staining with fluorescent phalloidin.

We assigned embryos to two broad age categories: early embryonic stages including comma to 1.5-fold, and late embryonic stages including 2-fold and older. Considering the elapsed 30 min temperature shift, we expected the younger embryos would normally have undergone initial polarization of F-actin and myosin during the temperature shift. We observed, however, a subset of early embryos lacked polarized muscle components (Fig. 4 A). Whereas a majority (∼ 80%) of wild-type, *fhod-1(Δ)*, or *cyk-1(ts)* early embryos had polarized BWMs, only 35% of *fhod-1(Δ); cyk-1(ts)* early embryos did so, the remainder being nonpolarized (Fig. 4 B). Surprisingly, embryos maintained at permissive temperature had similar percentages of abnormal BWM (Fig. S3 A), suggesting these defects were not due to an acute loss of CYK-1 function, but again consistent with *cyk-1(ts)* being partially non-functional at 16°C.

For late embryos of 2-fold stage and older, all embryos of all strains had sarcomeres, suggesting the early polarization defects of *fhod-1(Δ); cyk-1(ts)* double mutants are temporary. However, a fraction of embryos appeared abnormal in that F-actin-rich striations appeared frayed (Fig. 4 A). Again, proportion varied by genotype, with roughly 75% of wild type or *cyk-1(ts)* older embryos appearing normal, whereas only ∼ 10-15% *fhod-1(Δ)* or *fhod-1(Δ); cyk-1(ts)* embryos did so, the remaining 85-90% having frayed F-actin (Fig. 4 C, Fig. S3 B). Thus, absence of FHOD-1 perturbs sarcomere organization in late embryogenesis, but acute loss of CYK-1 does not significantly disrupt embryonic sarcomere assembly.

In order to verify the *cyk-1(ts)* was behaving as a loss of function allele under these test conditions, we also maintained populations of embryos at restrictive temperature overnight before fixation and staining with fluorescent phalloidin. We observed in some *fhod-1(*Δ*); cyk-1(ts)* L1 larvae detachment of the pharynx from the mouth (38% of animals at 26°C, versus 13% at 16°C, n = 100 animals per condition) (Fig. S4 A), a phenotype we have previously noted with longer-term absence of both formins in either *fhod-1(Δ); cyk-1(Δ)* mutants or after RNAi against *cyk-1* on *fhod-1(Δ)* mutants (Mi-Mi et al., 2012). In contrast, we never observed this in wild-type or *cyk-1(ts)* animals (n = 100 animals per strain per condition), and very rarely in *fhod-1(*Δ) animals (3% of animals at 16°C, 4% at 26°C, n = 100 animals per condition). Thus, short-term absence of CYK-1 does not prevent assembly of normal embryonic sarcomeres, but CYK-1 and FHOD-1 together contribute to normal pharyngeal development.

Unexpectedly, we also observed accumulation of F-actin in many nuclei of *cyk-1(ts)* and *fhod-1(*Δ*); cyk-1(ts)* embryos at permissive and restrictive conditions (Fig. 4 A, arrowheads, Fig. S4 B, C), but the significance of this is not clear. Overall, our data suggest FHOD-1 promotes proper F-actin organization in late embryonic BWM sarcomeres, whereas CYK-1 may be redundant with FHOD-1 in a non-essential role only during initial stages of sarcomerogenesis.

### FHOD-1 promotes BWM cell growth and proper DB organization in a cell-autonomous manner, whereas CYK-1 does not

Despite the fact that *cyk-1(*Δ) mutants exhibit significant BWM cell size and DB defects, our results here suggested CYK-1 is dispensable for embryonic sarcomere assembly, and makes minimal contributions to post-embryonic BWM development. One potential explanation for these discrepancies is that CYK-1 might promote BWM development indirectly, by playing a role in some other tissue that indirectly affects BWM. It also remained formally possible that FHOD-1 could also work in an indirect manner, as we had only examined constitutive *fhod-1(*Δ) mutants. Thus, we tested whether re-expression of FHOD-1 or CYK-1 could support BWM development in a muscle cell autonomous manner in *fhod-1(Δ)* and *cyk-1(Δ)* mutants, as would be expected if either directly contributes to sarcomere assembly.

We had previously demonstrated that full-length *fhod-1* genomic sequence with an encoded C-terminal GFP tag (denoted *fhod-1::gfp*) will partially restore BWM growth in *fhod-1(*Δ) mutants when integrated into an exogenous genomic site (Mi-Mi et al., 2012). We had also observed genomic integration of *cyk-1::gfp* derived from genomic full length *cyk-1* sequence rescues normal body size and partially rescues fertility of homozygous *cyk-1(*Δ) animals (Mi-Mi et al., 2012). With phalloidin stain of age-matched adult wild type, *cyk-1(*Δ), and transgene-rescued *cyk-1(*Δ*); cyk-1::gfp* animals, we confirmed exogenous *cyk-1* also restores normal BWM growth to *cyk-1(*Δ) mutants (Fig. S5).

To test for whether such BWM rescue occurs in a cell autonomous manner, we took advantage of the ability of *C. elegans* to host extrachromosomal arrays (ECAs) that tend to be inherited in a mosaic manner (Mello, Kramer, Stinchcomb, & Ambros, 1991). When plasmids are microinjected into the worm gonad, they are linearized and concatenated into ECAs that can be inherited through multiple generations of progeny, but often variably among cells within an embryo to produce mosaic animals. To rescue BWM defects, we microinjected plasmids bearing genomic sequences of full-length *fhod-1* or *cyk-1*. To avoid any potential interfering effects of tags, we utilized the original untagged formin genes from which the GFP-tagged versions had been created (Mi-Mi et al., 2012). To facilitate identifying transgenic BWM cells in transformed animals, we also co-injected a plasmid encoding free GFP expressed from the muscle-specific *myo-3* promoter. Concatenation of these plasmids into a single ECA in each worm strain ensured BWM cells expressing free GFP had also inherited the respective formin gene. As controls, animals were also injected with a mixture of non-formin DNA and the GFP*-* encoding plasmid.

For *fhod-1*, homozygous *fhod-1(*Δ) worms were microinjected and progeny were screened for mosaic expression of GFP (Fig. 5 A). For analysis, we selected two independent transformant lines derived from co-injection of *fhod-1(+)* with *gfp*, and two controls lines from co-injection of non-formin DNA with *gfp*. For *cyk-1*, we similarly injected *cyk-1(+)* or non-formin DNA, together with *gfp*-coding plasmid, but this was done into heterozygous *cyk-1*(Δ)/(+) worms to avoid the sterility of *cyk-1(*Δ) homozygotes.

**Figure 5.**
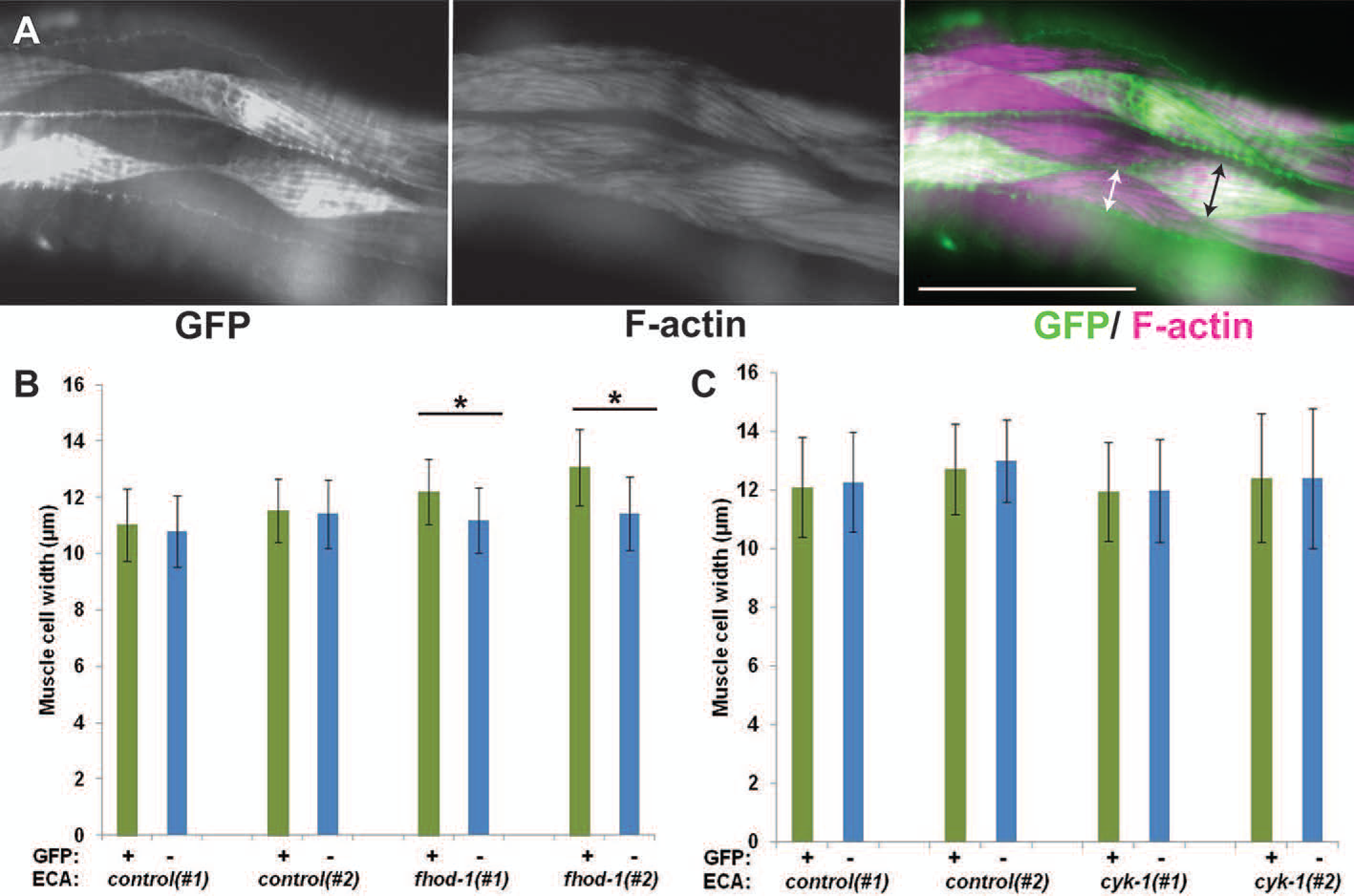
FHOD-1 plays a muscle cell autonomous role in promoting BWM cell growth whereas CYK-1 does not. Homozygous *fhod-1(Δ)* or heterozygous *cyk-1(Δ)/(+)* transgenic lines were selected for mosaic inheritance of extrachromosomal arrays (ECAs) containing wild type *fhod-1* or *cyk-1*, respectively, along with a gene encoding muscle-expressed GFP. Similar transgenic lines were selected for mosaic inheritance of ECAs constructed from non-formin DNA and the GFP marker. Two independently isolated lines of each genotype were selected for analysis. (A) Fluorescent phalloidin stain of a worm with mosaic expression of GFP allows identification of transgenic and non-transgenic BWM cells. Scale bar, 50 µm. Comparison of BWM cell widths of GFP-positive transgenic (black arrow) and GFP-negative non-transgenic (white arrow) cells allowed determination of cell autonomous effects of ECAs on muscle cell growth, as quantified in (B) and (C). (B) For *fhod-1(*Δ) strains bearing ECAs without *fhod-1*, designated here *control(#1)* and *control(#2)*, there was no significant effect of ECA presence on muscle cell size. Conversely, for *fhod-1(*Δ) strains bearing ECAs with *fhod-1*, designated *fhod-1(#1)* and *fhod-1(#2)*, transgenic BWM cells (green bars) were significantly wider than non-transgenic BWM cells in the same animals (blue bars). (C) For analysis of *cyk-1*, homozygous *cyk-1(Δ)* progeny of ECA-bearing *cyk-1(Δ)/(+)* parents were examined. Unlike for *fhod-1*, ECAs with *cyk-1* had no significant effect in *cyk-1(*Δ) animals on widths of transgenic BWM cells (green bars) compared to non-transgenic cells (blue bars). Graphs show the averages of the means of three experiments (n = 40 transgenic and 40 non-transgenic muscle cell widths measured in every experiment). Error bars indicate standandard error of the mean. (*) indicates p < 0.01. Differences in all other pairwise comparisons were not statistically significant (p > 0.05).

Synchronized adult mosaic worms were stained with phalloidin, and widths of individual adjacent GFP-positive (transgenic) muscle cells and GFP-negative (non-transgenic) muscle cells were measured (Fig. 5 A). As expected, there were no significant differences between GFP-positive and GFP-negative cells of control strains in either background (Fig. 5 B, C). However, in *fhod-1(*Δ) mutants with mosaic *fhod-1(+)* expression, *fhod-1-*bearing GFP-positive muscle cells were significantly wider than non-GFP neighbors (Fig. 5 B), suggesting FHOD-1 plays a cell autonomous role in promoting muscle cell growth. To test whether *cyk-1(+)* similarly promotes muscle cell growth, we examined BWM cells in homozygous *cyk-1(*Δ) progeny of heterozygous *cyk-1(*Δ*)/(+)* parents, which we identified by the presence of the *cyk-1(*Δ) phenotypes of protruding vulva and/or absence of embryos from the gonad (Mi-Mi et al., 2012). Consistently, widths of GFP-positive muscle cells expressing *cyk-1(+)* and GFP-negative cells not expressing *cyk-1(+)* were the same (Fig. 5 C), suggesting CYK-1 does not promote BWM cell growth in a cell autonomous manner.

Considering these results, we wanted to determine if FHOD-1 could also rescue DB organization in a cell autonomous manner. To test this, we performed immunostaining with anti-GFP and anti-ATN-1 on the same mosaic *fhod-1(*Δ) strains, and quantitatively analyzed the regularity of spacing of DBs along striations by performing fast Fourier transform (FFT) on ATN-1 immunostain intensity profiles along BWM striations. Consistent with a somewhat regular DB spacing in wild type animals, amplitude spectra for DBs in such animals showed a clustering of peaks near frequencies 0.8-1.0 µm^−1^, whereas irregularity of DB spacing in *fhod-1(*Δ) mutants correlated with spectra with no particular favored frequency (Fig. 6, S6). As expected, DBs of GFP-positive (transgenic) and GFP-negative (non-transgenic) BWM cells in *fhod-1(Δ)* strains transformed with non-formin DNA were irregular in shape and spacing, identical to *fhod-1(Δ)* mutants, and their amplitude spectra showed no favored frequency (Fig. 6, Fig. S6). In contrast, DBs in GFP-positive *fhod-1(+)*-expressing cells in the two mosaic rescued *fhod-1(*Δ) strains appeared regularly spaced, similar to wild-type, and their spectra peaks clustered near 0.8-1.0 µm^−1^ (Fig. 6, Fig. S6). Notably, neighboring GFP-negative (non-transgenic) cells in these same animals resembled those of non-transformed *fhod-1(Δ)* mutants, confirming FHOD-1 promotes proper DB organization in a cell autonomous manner.

**Figure 6.**
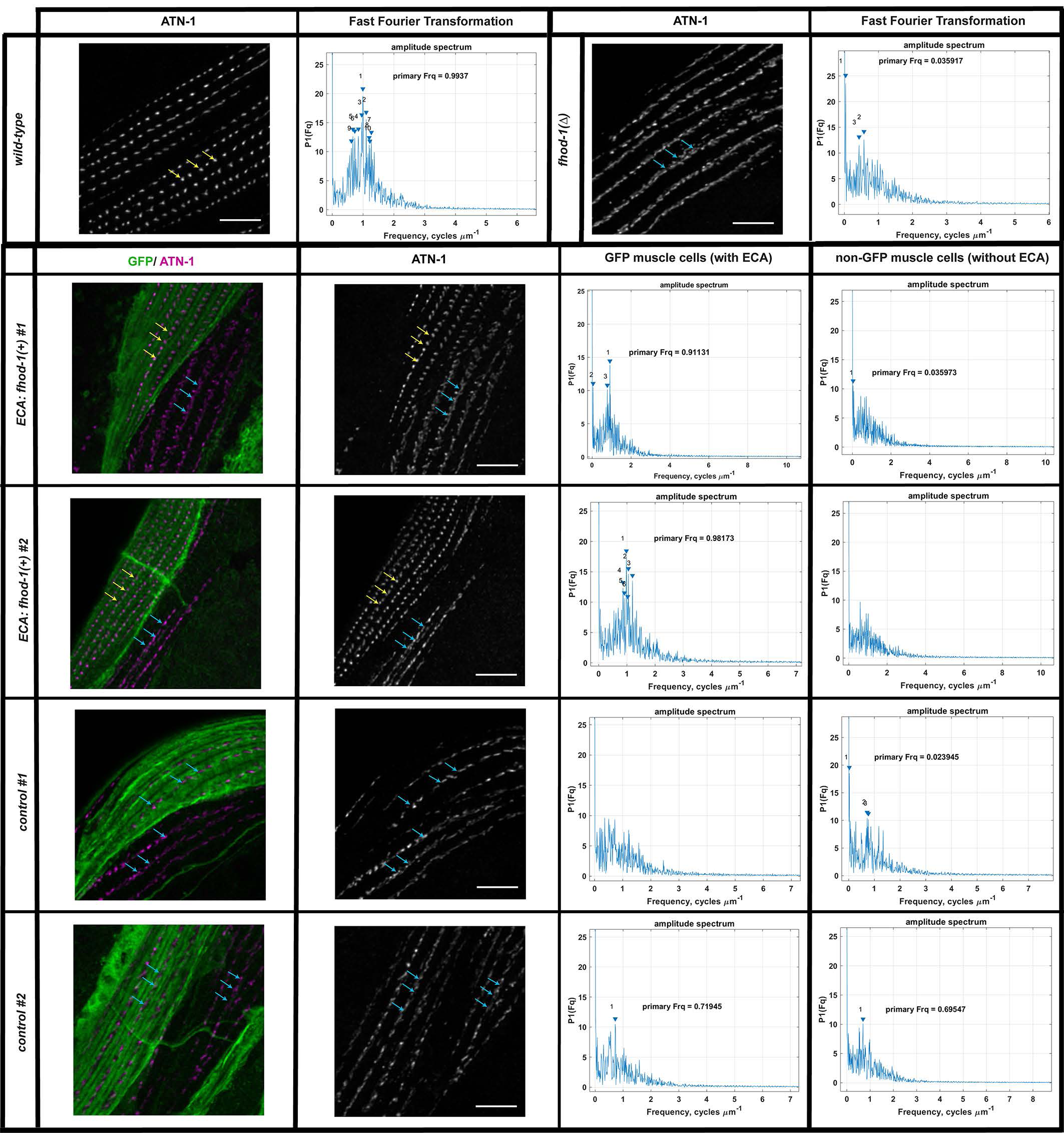
FHOD-1 plays a cell autonomous role in promoting proper DB morphology. Transformant lines of *fhod-1(*Δ) worms with mosaic inheritance of *fhod-1(+)-*containing or *fhod-1(+)-*lacking ECAs from Fig. 5 were immunostained for ATN-1 and GFP, along with wild type and non-transgenic *fhod-1(*Δ) animals. As shown in maximum intensity projections (MIP) of dorsal views from age-synchronized adult worms, ATN-1-positive DBs show a regular punctate distribution in wild type BWM, or in *fhod-1(*Δ) BWM cells that inherit a *fhod-1(+)*-containing ECA (yellow arrows), whereas DBs appear partially dispersed in *fhod-1(*Δ) BWM cells that inherit no ECA, or inherit an ECA lacking *fhod-1(+)* (blue arrows). DB spacing was quantitatively analyzed by performing fast Fourier transform (FFT) on concatenated intensity profiles of ATN-1 immunostain along approximately eight striations within single BWM cells (n = 10 animals per strain, one ECA-containing BWM cell and one non-ECA-containing BWM cell per animal for transgenic lines; one BWM cell for control strains). Amplitude spectra for wild type cells or *fhod-1(*Δ)cells that inherited a *fhod-1(+)-*containing ECA show clustering of peaks 0.8-1.0 µm^−1^, whereas spectra peaks did not cluster near any particular frequency for *fhod-1(*Δ) cells that inherited no ECA, or inherited an ECA lacking *fhod-1(+)*. Scale bars, 5 µm. Similar results were obtained for two additional replicate experiments (Fig. S6).

### FHOD-1 is enriched near sarcomeres in growing BWM cells throughout development, whereas CYK-1 does not localize to the contractile lattice in BWM cells

Our results so far indicated FHOD-1 contributes to BWM sarcomere organization throughout embryonic and postembryonic development. Previously, we had observed that FHOD-1 localizes in a diffuse pattern in BWM during late embryonic development after F-actin rich sarcomeres had assembled (older than 2-fold stage), and becomes enriched at BWM cell edges from mid larval development until early adulthood (Mi-Mi et al., 2012). Considering the apparent role we uncovered here for FHOD-1 during earlier embryonic muscle development, we re-examined its distribution in embryos and young larvae expressing a rescuing FHOD-1::GFP, after staining with fluorescently-labeled phalloidin to identify BWMs. Expanding on our previous results, we observed FHOD-1::GFP in BWMs of animals from embryonic 1.75 fold stage through L1 larva (Fig. 7). FHOD-1 localized diffusely in younger stage embryonic BWMs when sarcomeric components start to polarize and assemble into sarcomeres, but gradually appeared more punctuate at the edges of the F-actin-rich BWM cell contractile lattices from later embryogenesis into early larval development, when sarcomeres were maturing. Thus, FHOD-1 is present through all the stages of embryonic BWM development when sarcomere assembly is occurring.

**Figure 7.**
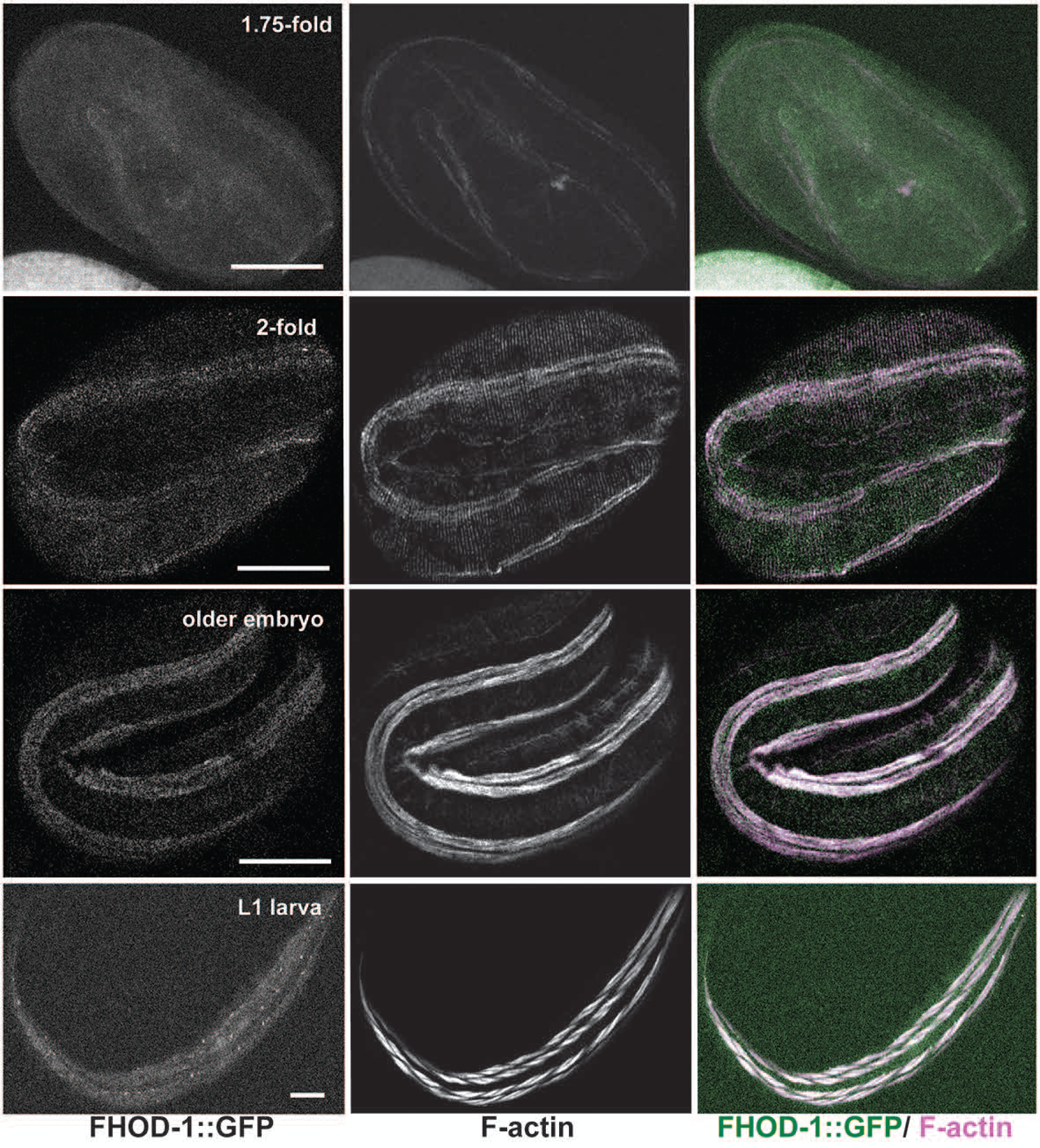
Distribution of FHOD-1::GFP in developing BWM. FHOD-1::GFP-expressing embryos of 1.75-fold, 2-fold, and older than 2-fold stages, and L1 larvae were stained with fluorescently-labeled phalloidin to visualize F-actin. FHOD-1::GFP localizes diffusely in early embryonic BWM, with gradual development of a punctuate appearance that becomes prominent in larval BWM. Scale bars, 20 µm.

Our results here also suggest CYK-1 plays no direct role in sarcomere formation, but the strong BWM defects in *cyk-1(Δ)* mutants reflect indirect contributions. This conclusion was surprising in light of previous evidence CYK-1 localizes to DBs, and thus we also re-examined CYK-1 localization. We previously observed CYK-1::GFP, when expressed from multiple copies of a transgene on an ECA, was present along striations in puncta that resembled DBs (Mi-Mi et al., 2012). However, that transgenic worm line lost the unstable ECA before we were able to compare the CYK-1::GFP puncta to a bona fide DB marker. To avoid the issue of ECA instability, we examined GFP fluorescence in a strain in which the endogenous *cyk-1* locus had been tagged with *gfp* using CRISPR-Cas9, resulting in a functional *cyk-1::gfp* (Davies et al., 2018). We observed CYK-1:: GFP localized in the germline similar to endogenous CYK-1 (Severson, Baillie, & Bowerman, 2002), but we did not observe localization to any particular structure in BWM, including DBs (Fig. S7 A, B). We also examined a strain expressing functional CYK-1::GFP from an ECA (Shaye & Greenwald, 2016), and observed punctate structures in BWM, but these were not positioned along the muscle I-bands or with any regularity, indicating these were not DBs (Fig. S7 C). Rather, these might represent aggregates that arose due to CYK-1::GFP over-expression from the ECA.

Considering these negative results, we tested the specificity of our previous anti-CYK-1 immunostain of DBs. For this, we performed *cyk-1(RNAi)* for 5 days on an RNAi-sensitive *cyk-1(+)* strain, as well as on the strain in which endogenous *cyk-1* had been tagged with *gfp*. Consistent with efficient knockdown, *cyk-1(RNAi)-*treated worms were sterile, and western blot analysis showed anti-CYK-1-reactive bands close to predicted molecular weights for CYK-1 (arrows) or CYK-1::GFP (arrowheads) were eliminated from *cyk-1(RNAi)*-treated animals (Fig. 8 A). Further confirming the efficacy of knockdown and specificity of anti-CYK-1 stain of the germline, immunostain was present in the germline in control animals (arrowheads) but not *cyk-1(RNAi)* animals (Fig. 8 B). In contrast, immunostain of adult worms with anti-CYK-1 and anti-MYO-3 (as counter-stain to visualize BWM, not shown), showed no difference between *cyk-1(RNAi)-*treated animals and controls (Fig. 8 C). Thus, BWM stain from anti-CYK-1 is likely non-specific, suggesting CYK-1 does not localize to DBs or any other sarcomeric structure in BWM, consistent with this formin playing no direct role in promoting sarcomere formation.

**Figure 8.**
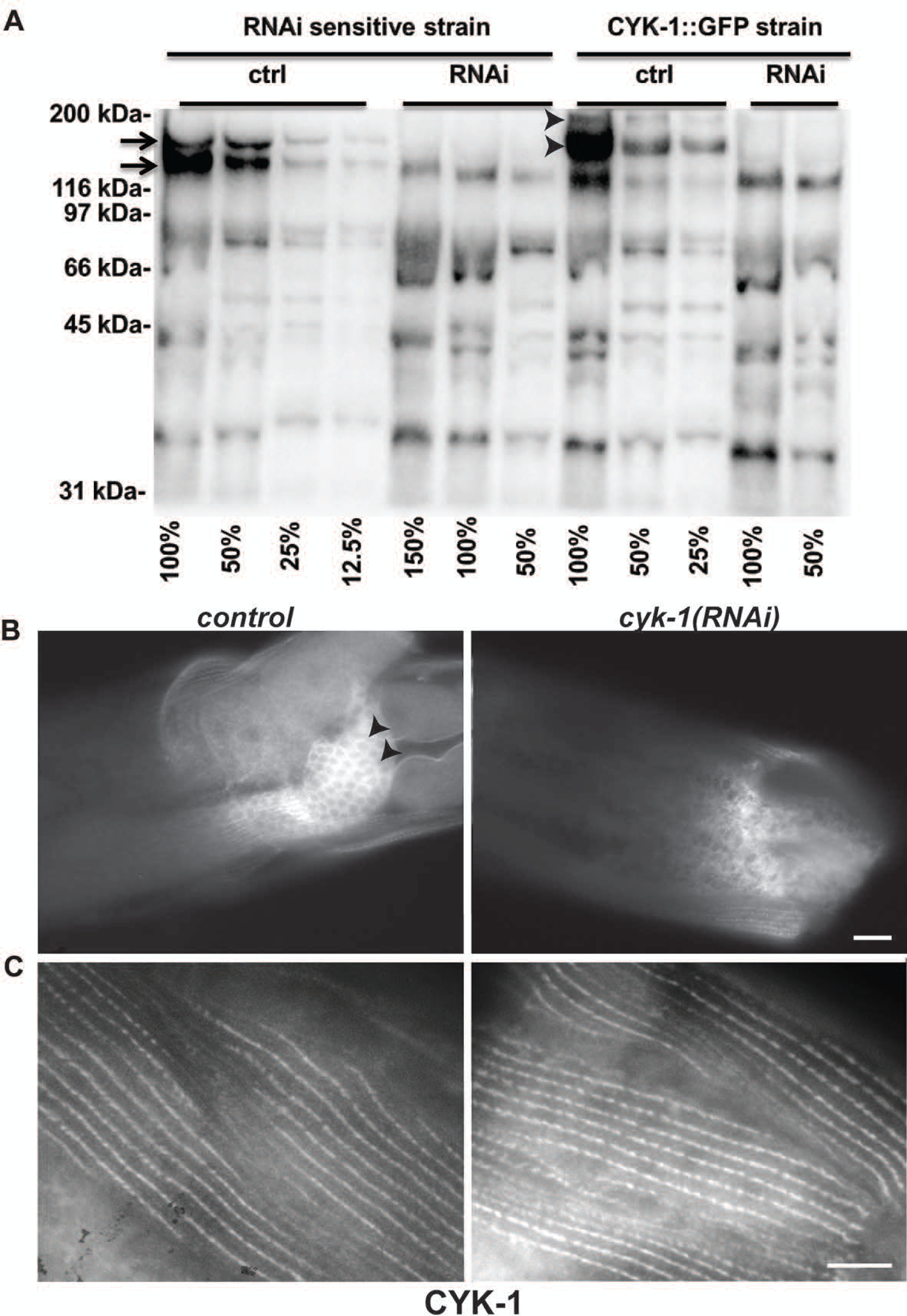
*cyk-1(RNAi)* does not alter DB-associated anti-CYK-1 immunostain. (A) Anti-CYK-1 western blot of dilutions (expressed as percentages) of adult worm extracts from RNAi sensitive *cyk-1(+)* animals, and animals in which endogenous *cyk-1* was tagged with GFP, treated with either control or *cyk-1(RNAi)*. Sample loads were approximately normalized based on previous whole lane protein determined by Coomassie Brilliant Blue staining. Bands of approximately 160 kDa and 170 kDa (arrows) for CYK-1 were lost from extracts of worms treated for *cyk-1(RNAi)* for 5 days, as were bands of approximately 187 kDa and 197 kDa (arrowheads) for CYK-1::GFP-expressing worm extracts. Non-specific bands recognized by this polyclonal anti-CYK-1 were not affected. (B) Anti-CYK-1 immunostaining shows CYK-1 localizes to the germline in control animals (arrowheads) and this localization is lost after *cyk-1(RNAi)*, but (C) anti-CYK-1 stain of DBs is similar between control and *cyk-1(RNAi)* animals, suggesting DB-associated stain is non-specific. Scale bars, 10 µm.

## Discussion

Formin contributions to sarcomere assembly during striated muscle development have been documented across various model systems, but the precise roles they play in this are unclear (Sanger et al., 2017). The striated body wall muscle (BWM) of *C. elegans* is an ideal system to test the roles of formins due to the worm’s simplicity in formin complement, which avoids potential redundancy due to the multiple isoforms of each formin family in vertebrates. Despite this simplicity, there is close similarity between sarcomeres in BWM and those in vertebrate cardiac muscle (Benian & Epstein, 2011). One compelling hypothesis is that formins could initiate thin filament assembly owing to the tendency of many formins to assemble long, unbranched actin filaments associated with tropomyosin.

We aimed to test that hypothesis using *C. elegans*, starting from our initial observations that simultaneous absence of actin assembly by two worm formins, FHOD-1 and CYK-1, profoundly stunts BWM development (Mi-Mi et al., 2012). Worms simultaneously bearing deletion alleles *fhod-1(*Δ*); cyk-1(*Δ) produce small BWM cells with a highly reduced number of sarcomeres, a phenotype that might be considered consistent with formins initiating thin filament assembly. However, the essential nature of CYK-1 made necessary maternal rescue of that particular *cyk-1(*Δ) allele, complicating this earlier analysis. Recent isolation of a conditional *cyk-1(ts)* (Davies et al., 2014) provided means to more fully eliminate CYK-1 activity after the critical window during embryogenesis had passed, without the complication of maternal inheritance.

Contrary to our expectations, analysis of *cyk-1(ts)* mutants suggested CYK-1 plays, at best, a very minor role in BWM growth during larval development. Muscle cell growth is driven by the assembly of additional sarcomeres, the majority of which occurs during larval development. Despite this, maintenance of developing *cyk-1(ts)* mutant larvae at a temperature where CYK-1 is non-functional caused almost no change in muscle cell growth, particularly when controlled for effects on overall body growth (Fig. 1). Even when *cyk-1(ts)* was paired with *fhod-1(*Δ), rates of muscle cell growth were essentially identical to those for larvae bearing just *fhod-1(*Δ).

Similarly, BWM Z-line defects of *cyk-1(*Δ) mutants were not recapitulated in *cyk-1(ts)* animals. That is, in electron micrographs, wild type dense bodies (DBs, the BWM Z-line analogs) appear as electron-dense finger-like projections, but many DBs in *cyk-1(Δ)* mutants appear deficient in electron-dense material (Mi-Mi & Pruyne, 2015). Correspondingly, DBs of wild-type animals appear as ATN-1 (α-actinin)-rich puncta by immunostain, but many regions of *cyk-1(Δ)* BWM lack apparent ATN-1-stained DBs (Fig. 2). Conversely, DBs of *cyk-1(ts)* mutants grown at a restrictive temperature throughout larval development appear normal (Fig. 2), again despite most DBs having assembled during the restrictive period. FHOD-1 absence also affects DB morphology (discussed below), but pairing of *cyk-1(ts)* with *fhod-1(*Δ) did not produce a more severe phenotype than that of *fhod-1(*Δ), alone (Fig. 2).

We suspected these differences in phenotypes might be due to differences in the timing of CYK-1 loss between *cyk-1(Δ)* and *cyk-1(ts)*. That is, through temperature control we restricted loss in *cyk-1(ts)* mutants to post-embryonic development, whereas CYK-1 loss presumably occurs earlier in *cyk-1(Δ)* mutants with depletion of maternal product. The more severe *cyk-1(*Δ) effects might therefore imply that CYK-1 contributes to BWM formation earlier in development. However, using the *cyk-1(ts)* we were unable to find evidence for a significant CYK-1 role in embryonic BWM sarcomere formation. With acute removal of CYK-1 activity, embryos of *fhod-1(*Δ*); cyk-1(ts)* double mutants did exhibit a defect in initial accumulation of F-actin and myosin at the start of sarcomere formation (Fig. 4), suggesting FHOD-1 and CYK-1 may redundantly promote early polarization of muscle components. However, we observed this phenotype at similar frequencies at the permissive temperature (Fig. S3), suggesting it is not due to acute removal of formin activity during sarcomerogenesis. Moreover, as sarcomeres are present in the vast majority of two-fold stage *fhod-1(*Δ*); cyk-1(ts)* embryos, we reasoned this early polarization defect is probably temporary.

One caveat to using the *cyk-1(ts)* is it remained possible the temperature-sensitive CYK-1 protein was not actually losing function in BWM. Thus, we used the alternative approach of examining the strong BWM defects of *cyk-1(*Δ) mutants. We confirmed genomic integration of a *cyk-1* transgene fully rescues BWM defects of *cyk-1(*Δ) animals (Fig. S5), thereby demonstrating those defects are specific to loss of CYK-1. However, when a similar *cyk-1* transgene was inherited in a mosaic pattern, we failed to observe cell autonomous rescue of BWM cell growth by the transgene (Fig. 5 C).

It was surprising that we also saw no evidence of any rescue at all of BWM cell growth in these mosaic *cyk-1-*expressing animals. That is, if *cyk-1* promotes BWM development in a non-cell autonomous manner, we might have expected all BWM cells of mosaic *cyk-1* animals to be larger than BWM cells of control animals lacking the transgene. However, we observed no difference in BWM cell size between those strains (Fig. 5 C, control versus *cyk-1*). Why might a mosaic pattern of expression not support the function, while a genome-integrated transgene would? Possible explanations include *cyk-1* might be required simultaneously in many cells, or in a tissue where expression of non-integrated transgenes is weak, such as in the germline (Mello et al., 1991).

Our overall results, together with inability to detect CYK-1 in BWM sarcomeres (Fig. 8, Fig. S7), suggest CYK-1 plays an indirect role in BWM development, likely functioning in a tissue or tissues other than BWM, and this likely occurs during embryogenesis. Consistent with CYK-1 functioning in tissues other than BWM, *cyk-1(*Δ) or *cyk-1(RNAi)-*treated animals exhibit a range of defects in non-muscle tissues, including germline, epidermis, intestine, and excretory canal (Swan et al., 1998; Mi-Mi et al., 2012; Shaye & Greenwald, 2016; Gong et al., 2018). We also observed that in embryos bearing *cyk-1(ts)* there is widespread accumulation of F-actin in nuclei (Fig. 4 A, Fig. S4 B, C), showing loss of CYK-1 has widespread effects throughout the body even during very early development.

In contrast, FHOD-1 seems to promote BWM development and Z-line organization directly. FHOD-1 is initially diffuse in embryonic BWM cells, when sarcomeric components begin to accumulate at the cell membrane. Once the initial sarcomeres have formed, the formin appears as puncta at BWM cell edges, and more diffusely along sarcomere I bands (Fig. 7), remaining so until the end of BWM growth in adulthood, when localized formin is no longer detected (Mi-Mi et al., 2012). In embryos, lack of FHOD-1 results frayed-appearing F-actin in the newly formed sarcomeres (Fig. 4, Fig. S4), suggesting a possible partial defect in thin filament anchorage and organization. In larvae lacking FHOD-1, additional sarcomere assembly in BWM cells is slowed, resulting in smaller BWM cells (Fig. 1, Fig. S1) (Mi-Mi et al., 2012). Contrasting CYK-1, mosaic expression of FHOD-1 in *fhod-1(*Δ) mutants promotes BWM cell growth in a cell autonomous manner (Fig. 5). Additionally, DBs are of irregular size and spacing in *fhod-1(*Δ) adults (Mi-Mi et al., 2012), but mosaic FHOD-1 expression also rescues DB organization in a cell autonomous manner (Fig. 6, Fig. S6). These results suggest the ability of FHOD-1 to promote BWM cell growth and proper DB organization are direct effects that may be functionally linked.

We have found no phenotypic evidence of any other formin contributing to BWM development in worms, either from individual formin gene mutations, or from pairing of *fhod-1(*Δ) with mutations in each of the remaining non-*cyk-1* formins (Fig. S2) (Mi-Mi et al., 2012). One caveat to this would be possibility of a higher degree of redundancy among formins for BWM development, with simultaneous elimination of more (or all) worm formins resulting in a stronger phenotype, such as disruption of thin filament assembly. However, absence of even hints of synthetic defects in BWM development between *fhod-1(*Δ) and other non-*cyk-1* formin mutations makes this unlikely. Thus, we suggest *fhod-1(*Δ) BWM cells lack any formin activity that contributes directly to sarcomere formation. And as *fhod-1(*Δ) BWM cells contain abundant thin filaments, we think it unlikely that thin filament assembly in *C. elegans* BWM requires formins. It is possible this differs in vertebrate or fly muscle, and that formins assemble their thin filaments. However, vertebrate muscle contains other non-formin actin nucleating factors that might initiate thin filament assembly, including leiomodins (Chereau et al., 2008; Yuen et al., 2014; Boczkowska, Rebowski, Kremneva, Lappalainen, & Dominguez, 2015) and a complex of N-WASP with nebulin (Takano et al., 2010). Worms lack unambiguous leiomodin and nebulin homologs, but they host related proteins, and other unknown actin nucleation factors could initiate thin filament assembly.

It is unclear why worm muscle is so resilient to formin loss compared to other organisms. Loss of DAAM-family formins significantly perturbs sarcomere organization in mammalian and insect muscle (Molnar et al., 2014; Ajima et al., 2015), while worm DAAM-1 appears dispensable (Fig. S2). Absence of FHOD-1, the only *C. elegans* representative of FHOD-family, results in muscle defects much milder than the well-studied impacts of FHOD-family members in other systems (Taniguchi et al., 2009; Iskratsch et al., 2010; Kan-O et al., 2012; Shwartz et al., 2016; Ushijima et al., 2018; Fenix et al., 2018). For example, where *fhod-1(*Δ) worms exhibit reduced sarcomere assembly and partially defective Z-lines/DBs, knockdown of FHOD3 in human induced pluripotent stem cell-derived cardiomyocytes blocks the maturation of stress fiber-like structures into sarcomere-containing myofibrils (Fenix et al., 2018), phenotypes that appears to recapitulate those of the *fhod3*^*-/-*^ mouse heart (Kan-O et al., 2012). Moreover, where *fhod-1(*Δ) BWM is largely functional and mutant worms are fully viable, mice lacking FHOD3 die during embryonic development due to heart failure (Kan-O et al., 2012).

One possible explanation for why worm BWM is resistant to formin loss is the unique architecture of its contractile machinery. Sarcomeres in mammalian and *Drosophila* striated muscles organize into myofibrils, most of which are suspended in the cytoplasm away from the plasma membrane. Conversely, all BWM sarcomeres are directly anchored to the plasma membrane and the underlying extracellular matrix through integrin-based adhesions. This likely provides significant mechanical reinforcement. This might, for example, permit DBs in worm muscle to tolerate structural defects that would be catastrophic for Z-lines in myofibrils. The mechanisms for precisely how mammalian FHOD3 or worm FHOD-1 promotes sarcomere formation are not understood, but the relative resilience of worm muscle to loss of its FHOD-family formin may prove to be an advantage in dissecting details of this process.

## Materials and Methods

### Worm strains and growth conditions

Worms were maintained and grown using standard protocols (Brenner, 1974) at 20°C, except for experiments involving temperature-sensitive worms, for which growth was at 16°C prior to temperature shifts. Age-synchronized populations were obtained in one of two ways. By one method, adult worms were treated with 1:2 ratio 5 mM NaOH to reagent grade bleach to liberate embryos, which were then washed with M9 buffer, and allowed to develop until the proportion at the two-fold stage had peaked (Fig. 3, 4, S3, S4), or until embryos developed into starvation-arrested L1 stage larvae (Fig. 1, 2, 7, 8, S1, S4). Alternatively, adult animals were allowed to lay eggs on plates for about ∼ 8 hrs and then were removed, resulting in semi-synchronized progeny (Fig. 5, 6, S2, S5, S6, S7).

For long temperature shift experiments (Fig. 1, 2, S1), starvation-arrested L1 stage larvae were transferred to plates with food (*Escherichia coli* OP50) at permissive (16°C) or restrictive (26°C) temperature for 0, 12, 24, 36, 48, 60, or 72 hrs, before being prepared for fluorescence microscopy. For short temperature shift experiments (Fig. 4, S3, S4), embryos in M9 were transferred to 16°C or 26°C water baths for 30 min, before preparation for fluorescence microscopy.

For complete genotypes of strains used in this study, see Table 1. N2 (wild type Bristol worms) and RW1596 were supplied by *Caenorhabditis* Genetics Center (University of Minnesota, Minneapolis, MN). JCC389 and JCC955 were gifts from Julie Canman (Columbia University, New York, NY; Davis et al., 2014; Davies et al., 2018). XA8001, DWP10, DWP22, and DWP28 were obtained previously (Mi-Mi et al., 2012). DWP68 was obtained by crossing JCC389 with XA8001. DWP156, DWP160, and DWP161 were obtained by crossing XA8001, JCC389 and DWP68, respectively, into the RW1596 background. GS7933 was a gift from Daniel Shaye (University of Illinois at Chicago, Chicago, IL; Shaye & Greenwald, 2016). DWP153 and DWP154 were created by crossing worms bearing the balancer *mT1[dpy-10(e128)]* with DWP8 [*cyk-1(ok2300)/+*III] and DWP9 *fhod-1(tm2363)*; *cyk-1(ok2300)/+*III], respectively (Mi-Mi et al., 2012).

**Table 1.**
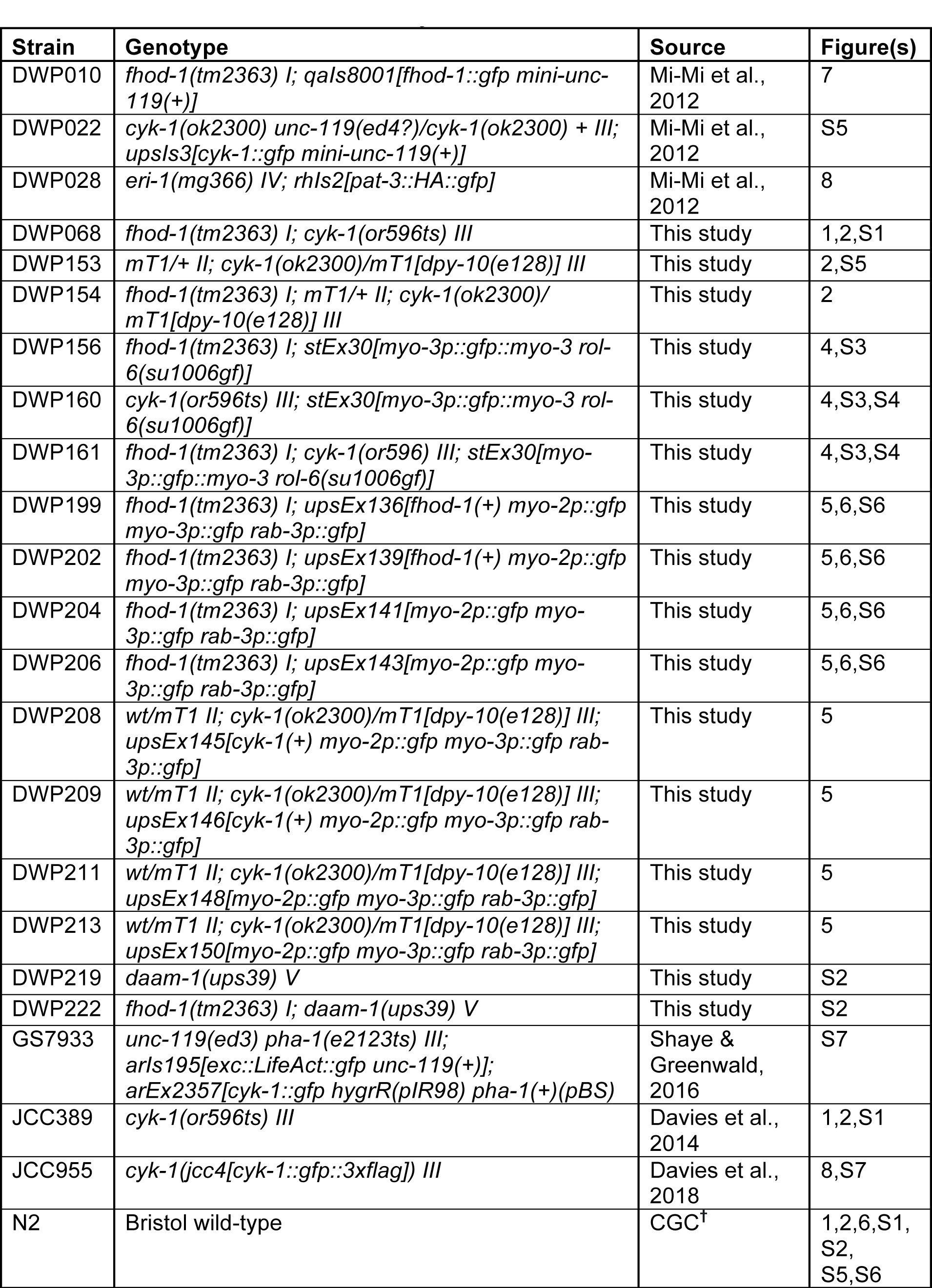
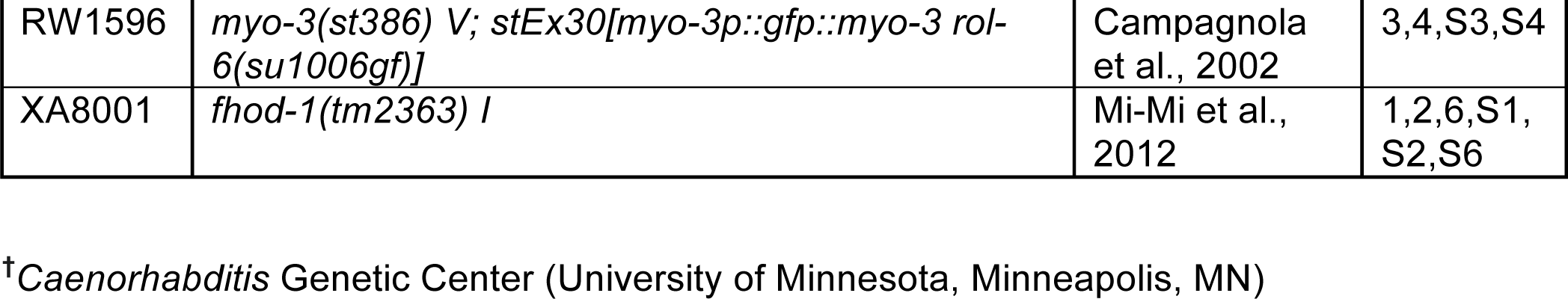
Worm strains used in this study. Strain names with respective genotypes, figure(s) in which they appear, and source.

| Strain | Genotype | Source | Figure(s) |
| --- | --- | --- | --- |
| DWP010 | *fhod-1(tm2363) I; qals8001[fhod-1::gfp mini-unc-119(+)]* | Mi-Mi et al., 2012 | 7 |
| DWP022 | *cyk-1(ok2300) unc-119(ed4?)/cyk-1(ok2300) + III; upsls3[cyk-1::gfp mini-unc-119(+)]* | Mi-Mi et al., 2012 | S5 |
| DWP028 | *eri-1(mg366) IV; rhls2[pat-3::HA::gfp]* | Mi-Mi et al., 2012 | 8 |
| DWP068 | *fhod-1(tm2363) I; cyk-1(or596ts) III* | This study | 1,2,S1 |
| DWP153 | *mT1/+ II; cyk-1(ok2300)/mT1[dpy-10(e128)] III* | This study | 2,S5 |
| DWP154 | *fhod-1(tm2363) I; mT1/+ II; cyk-1(ok2300)/mT1[dpy-10(e128)] III* | This study | 2 |
| DWP156 | *fhod-1(tm2363) I; stEx30[myo-3p::gfp::myo-3 rol-6(su1006gf)]* | This study | 4,S3 |
| DWP160 | *cyk-1(or596ts) III; stEx30[myo-3p::gfp::myo-3 rol-6(su1006gf)]* | This study | 4,S3,S4 |
| DWP161 | *fhod-1(tm2363) I; cyk-1(or596) III; stEx30[myo-3p::gfp::myo-3 rol-6(su1006gf)]* | This study | 4,S3,S4 |
| DWP199 | *fhod-1(tm2363) I; upsEx136[fhod-1(+) myo-2p::gfp myo-3p::gfp rab-3p::gfp]* | This study | 5,6,S6 |
| DWP202 | *fhod-1(tm2363) I; upsEx139[fhod-1(+) myo-2p::gfp myo-3p::gfp rab-3p::gfp]* | This study | 5,6,S6 |
| DWP204 | *fhod-1(tm2363) I; upsEx141[myo-2p::gfp myo-3p::gfp rab-3p::gfp]* | This study | 5,6,S6 |
| DWP206 | *fhod-1(tm2363) I; upsEx143[myo-2p::gfp myo-3p::gfp rab-3p::gfp]* | This study | 5,6,S6 |
| DWP208 | *wt/mT1 II; cyk-1(ok2300)/mT1[dpy-10(e128)] III; upsEx145[cyk-1(+) myo-2p::gfp myo-3p::gfp rab-3p::gfp]* | This study | 5 |
| DWP209 | *wt/mT1 II; cyk-1(ok2300)/mT1[dpy-10(e128)] III; upsEx146[cyk-1(+) myo-2p::gfp myo-3p::gfp rab-3p::gfp]* | This study | 5 |
| DWP211 | *wt/mT1 II; cyk-1(ok2300)/mT1[dpy-10(e128)] III; upsEx148[myo-2p::gfp myo-3p::gfp rab-3p::gfp]* | This study | 5 |
| DWP213 | *wt/mT1 II; cyk-1(ok2300)/mT1[dpy-10(e128)] III; upsEx150[myo-2p::gfp myo-3p::gfp rab-3p::gfp]* | This study | 5 |
| DWP219 | *daam-1(ups39) V* | This study | S2 |
| DWP222 | *fhod-1(tm2363) I; daam-1(ups39) V* | This study | S2 |
| GS7933 | *unc-119(ed3) pha-1(e2123ts) III; arls195[exc::LifeAct::gfp unc-119(+)]; arEx2357[cyk-1::gfp hygrR(plR98) pha-1(+)(pBS)]* | Shaye & Greenwald, 2016 | S7 |
| JCC389 | *cyk-1(or596ts) III* | Davies et al., 2014 | 1,2,S1 |
| JCC955 | *cyk-1(jcc4[cyk-1::gfp::3xflag]) III* | Davies et al., 2018 | 8,S7 |
| N2 | Bristol wild-type | CGC<sup>†</sup> | 1,2,6,S1, S2, S5,S6 |
| RW1596 | *myo-3(st386) V; stEx30[myo-3p::gfp::myo-3 rol-6(su1006gf)]* | Campagnola et al., 2002 | 3,4,S3,S4 |
| XA8001 | *fhod-1(tm2363) I* | Mi-Mi et al., 2012 | 1,2,6,S1,S2,S6 |

<sup>†</sup>*Caenorhabditis* Genetic Center (University of Minnesota, Minneapolis, MN)

Novel loss-of-function allele of *daam-1(ups39)* was generated using CRISPR/Cas9-mediated gene editing to target TAATTGACCCGAGACGCTAT near the start of an FH2-coding exon of *daam-1*. A plasmid to direct homologous recombination-guided repair was constructed through the following steps. By PCR, a fragment encoding nucleotides 22,974 – 24,147 of *daam-1* was amplified with appended upstream *KpnI* cloning site, and downstream *EcoRI* site, in-frame stop codon, *LoxP* site, and *SalI* cloning site (primers GGTACCGAGCGGATTGCAAAAGAGCTGGA and GTCGACATAACTTCGTATAATGTATGCTATACGAAGTTATTCAGAATTCGGTGATCTGGAAATGAAAGTTGTATAC). Similarly, nucleotides 24,198 – 25,339 of *daam-1* were amplified and appended with upstream *BamHI* cloning site and *LoxP* site, and downstream *NotI* cloning site (primers GGATCCATAACTTCGTATAGCATACATTATACGAAGTTATAACTGCACAATAATGCTCTCCA AGC and GCGGCCGCTCTACCCACCTCATACCTACACGC). These were sequentially cloned into pJKL702 (a gift from Jun Kelly Liu, Cornell University, Ithaca, NY) to flank its *unc-119* mini-gene. Using this homologous recombination-guided repair template, we generated and isolated transgenic lines from HT1593 [*unc-119(ed3)* III] animals, and excised the integrated *unc-119(+)* from the *daam-1* locus by expression of Cre recombinase, all as described by others (Dickinson & Goldstein, 2016). The resultant post-excision allele *daam-1(ups39)* encodes an in-frame stop codon near the start of the FH2-coding sequence, and a 1-nt frame shift due to the *LoxP* site, and is thus predicted encode a non-functional formin. DWP219 was generated by crossing *daam-1(ups39)* into an *unc-119(+)* background, using EcoRI-mediated cleavage as a diagnostic for *ups39*. DWP222 was created by crossing DWP219 with XA8001.

To generate worms with mosaic expression of *fhod-1*, young adult XA8001 hermaphrodites were microinjected with a mixture of 50 ng/µL pRS315-*fhod-1(+)* (Mi-Mi et al., 2012), 100 ng/µL *rab-3p::gfp* plasmid, 50 ng/µL *myo-3p::gfp* plasmid, and 25 ng/µL *myo-2p::gfp* plasmid. Two resulting strains, DWP199 and DWP202, were isolated from progeny of different injected parents for their expression of GFP in a mosaic pattern in BWM. Control strains DWP204 and DWP206, with mosaic transgene inheritance but lacking *fhod-1(+)*, were produced similarly, but replacing pRS315-*fhod-1(+)* with pRS315 (Sikorski & Hieter, 1989). To isolate animals with mosaic expression of *cyk-1(+)*, DWP153 worms were microinjected with 50 ng/µL pRS315-*cyk-1(+)* or pRS315, and 100 ng/µL *rab-3p::gfp* plasmid, 50 ng/µL *myo-3p::gfp* plasmid, and 25 ng/µL *myo-2p::gfp* plasmid. The resulting strains, DWP208 and DWP209 with transgenic *cyk-1(+)*, and control strains DWP211 and DWP213 without transgenic *cyk-1(+)*, were identified by selecting for progeny expressing GFP in a mosaic pattern in BWM, again from different injected parents.

### RNAi

RNAi-mediated knockdowns were performed by the standard feeding technique (Wang and Barr, 2005). Briefly, *E. coli* HT115 was transformed with control knockdown vector L4440 (Timmons & Fire, 1998) or *cyk-1* knockdown vector L4440-*cyk-1* (Mi-Mi et al., 2012), and grown overnight at 37°C in 2xYT with 12.5 µg/ml tetracycline, 100 µg/ml ampicillin. Cultures were then diluted 1:100 in 2xYT, grown 3 hrs 37°C, and induced 3-4 hrs with 0.4 mM IPTG at 37°C. Induced cultures were concentrated five-fold and seeded onto plates. Age-synchronized L1 worms were plated and inspected as adults after 3-4 days at 25°C. Adult worms were washed with M9 to remove progeny, and plated on freshly induced bacteria to continue RNAi knockdown for 5 days total treatment.

### Staining for fluorescence microscopy

F-actin stain was as previously described (Mi-Mi et al., 2012), and immunostain as previously described (Finney & Ruvkun, 1990), except for omission of spermidine-HCl from the initial buffer for both protocols, and threefold higher initial methanol concentration (75%) for immunostain. Monoclonal primary antibody MH35 (anti-ATN-1) generated by R.H. Waterston (Francis & Waterston, 1985) was a gift from Pamela Hoppe (Western Michigan University, Kalamazoo, MI), polyclonal rabbit anti-GFP was a gift from Anthony Bretscher (Cornell University, Ithaca, NY), polyclonal affinity-purified anti-CYK-1 (DPMSP1) was generated as previously described (Mi-Mi et al., 2012), mouse anti-GFP and secondary antibodies (Texas red-conjugated goat anti-rabbit and FITC-conjugated goat anti-mouse, or reverse fluorophore/species) were commercial (Rockland Immunochemicals, Pottstown, PA). Antibody dilutions were 1:200 DPMSP1, 1:10^4^ MH35, 1:10^3^ mouse anti-GFP, 1:200 rabbit anti-GFP, and 1:500 for secondary antibodies.

### Microscopy and image analysis

Wide field fluorescence images (Fig. 1, 4, 5, 8 B, C, S2, S4 A, B, S5, S7 A) were acquired using an Eclipse 90i Upright Research Microscope (Nikon, Tokyo, Japan) at room temperature (∼25°C) with a CFI Plan Apochromat 40x/NA 1.0 oil immersion objective, or a CFI Plan Apochromat violet-corrected 60x/NA 1.4 oil immersion objective, with a Cool-SNAP HA2 digital monochrome charge-coupled device camera (Photometrics, Tucson, AZ) driven by NIS-Elements AR acquisition and analysis software (version 3.1; Nikon, Tokyo, Japan). Confocal images (Fig. 2, 3, 6, 7, S4 C, S7 B, C) were obtained on an SP8 Laser Scanning Confocal Microscope (Leica, Wetzlar, Germany) driven by LAS X Software (version 2.2.0, build 4758; Leica, Wetzlar, Germany), and using an HCX Plan Apochromat 63x/NA 1.4 oil lambda objective. Maximum intensity projections and XZ/YZ cross-sections were generated from confocal stacks using LAS AF Software or ImageJ (version 2.0.0-rc-65/1.51g) (Schneider, Rasband, & Eliceiri, 2012). All images were linearly processed to enhance contrast and false-colored in Photoshop CS4 or Photoshop CC 2018 (Adobe, San Jose, CA).

BWM width and muscle cell width were measured based on phalloidin stain, and total body width was measured based on body autofluorescence in the FITC channel, as described previously (Mi-Mi et al., 2012) using NIS-Elements AR acquisition software. Embryonic developmental stages were visually identified based on embryo shape, and BWMs were identified by the presence of GFP::MYO-3 (see results, Fig. 3, 4). In animals with mosaic expression of soluble GFP in muscle cells, the FITC-channel was used to locate adjacent green and non-green muscle cells, followed by analysis of muscle cell width based on phalloidin stain, or analysis of DB organization based on anti-ATN-1 immunostain (Fig 5, 6, S6). To examine BWM cells in homozygous *cyk-1(*Δ) progeny of heterozygous *cyk-1(*Δ*)/(+)* parents (Fig. 5), such progeny were identified through the presence of protruding vulva and/or absence of embryos from the gonad (Mi-Mi et al., 2012). For RNAi experiments requiring examination of CYK-1 immunostain after *cyk-1(RNAi)* (Fig.8), we selected animals negative for anti-CYK-1 germline stain as verification of efficient knockdown of CYK-1. Fluorescence images for comparison between *cyk-1(RNAi)* and control (Fig. 8) were consistently acquired at 500 ms exposures. All quantitative analyses were performed while blinded to the strain genotype.

### Fast Fourier transform (FFT)

Anti-ATN-1 stain intensity values were obtained for approximately eight DB-containing striations in one GFP-positive (ECA-containing) muscle cell and one adjacent GFP-negative (ECA-lacking) muscle cell in each of ten mosaic animals of each strain (as seen in Fig. 5), using Freehand Line tool in ImageJ. The number of DBs per striation is relatively small, and their spacing is somewhat variable, which produces a relatively weak signal after FFT. To amplify this signal, intensity profiles of a given kind (e.g. profiles from all eight striations from one GFP-positive cell from each of ten animals of a given genotype) were concatenated. To ensure spacing was not disrupted by concatenation, intensity profiles were joined at peak maxima for ATN-1 (i.e. the last peak maximum of the intensity profile from the one cell/striation was joined to the first peak maximum of the intensity profile from the next cell/striation, and so on). FFT was performed and amplitude spectra obtained using MATLAB (R2019a Update 2). All analyses were performed while blinded to strain genotypes.

### Western blots

Whole worm lysates were obtained from age-synchronized young adult worms by washing worms off plates and separating from *E. coli* before concentrating in 1.7 mL tubes to a 1:1 worm to M9 slurry. Reducing sample buffer (2X) was directly added to samples before boiling 3 min, disruption 30 sec with tissue homogenizer (VWR International, Radnor, PA), boiling 3 min, and pelleting 15 sec. To break up genomic DNA, samples were pulled through an insulin syringe eight times before loading for SDS-PAGE. For purposes of normalization, samples were subject to preliminary SDS-PAGE and Coomassie brilliant blue stain. Images were acquired using a Bio-Rad ChemiDoc MP imager (Bio-Rad, Hercules, CA), and intensities of stain for total lanes were compared using Image Lab software (Bio-Rad, Hercules, CA). For western blot analysis, proteins in normalized samples were resolved by SDS-PAGE and transferred to nitrocellulose (Bio-Rad, Hercules, CA). Blots were blocked in 10% milk in TBST (50 mM Tris-HCl, pH 8.3, 150 mM NaCl, 0.3% Tween 20) and incubated with DPMSP1 primary antibody diluted 1:200 in TBST/1% milk, 5 hrs at room temperature. Blots were washed and incubated in goat anti-rabbit horseradish peroxidase-conjugated secondary antibody (Rockland Immunochemicals, Gilbertsville, PA) diluted 1:3000 in TBST/1% milk, 1 hr at room temperature, before washing and treating with enhanced chemiluminescence substrate (ThermoFisher Scientific, Waltham, MA). Images were acquired using a Bio-Rad ChemiDoc MP imager (Hercules, CA), and processed with Image Lab and Photoshop CS4.

### Statistical analyses

Numerical data in the text are expressed as mean ± standard error of the mean or one standard deviation, as defined in the text. Graphs were made in Excel:windows (version 14.7.2; Microsoft Corporation, Redmond, WA). For results where two groups are compared (Fig. 5), data were analyzed using a student t-test, with p < 0.05 considered statistically significant. For results from three or more groups, data were analyzed using Analysis of Variance, followed by a Least Significant Difference *post hoc* test. P-value < 0.05 was considered to be statistically significant.

## Acknowledgements

Thanks to Julie Canman for worm strains, Peter Calvert for help with FFT analysis, and WormBase and WormAtlas. Worm strains were obtained from the CGC, which is funded by NIH Office of Research Infrastructure Programs (P40 OD010440). This work was supported by NIAMS of the NIH under Award Number R01AR064760 to D.P. and Charles H. Revson Senior Fellowship in Biomedical Science to T.D.

---

**Figure S1.**
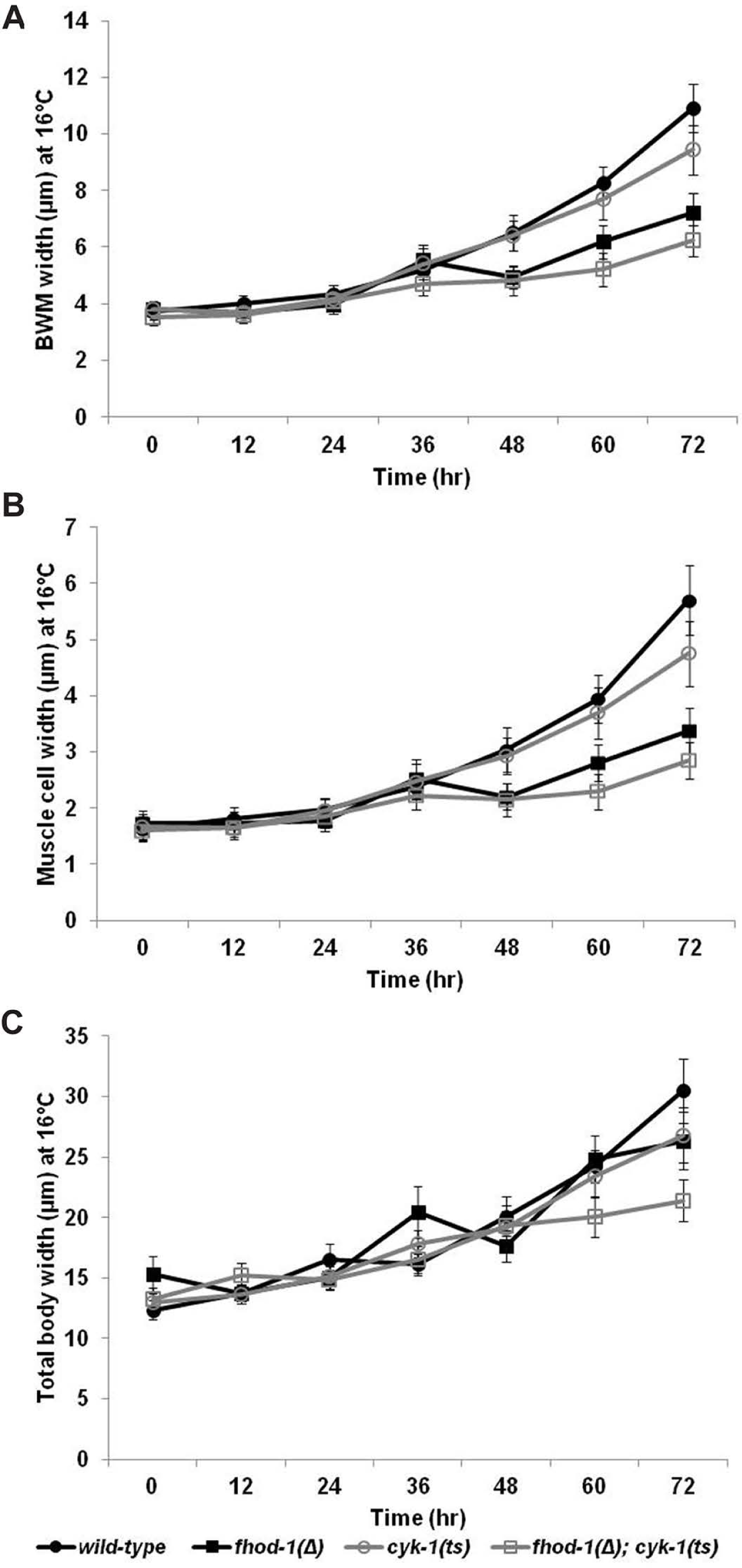
*cyk-1(ts)* mutants grown at permissive temperature. Embryos were hatched and grown at permissive temperature (16°C) for up to 72 hrs. Samples were collected every 12 hrs, stained with fluorescently-labeled phalloidin, and measured for (A) BWM width, (B) individual muscle cell width, and (C) total body width. We observed modestly reduced BWM, individual muscle cell, and total body widths in *cyk-1(ts)* mutants compared to wild-type at permissive temperatures. Graphs depict the average of the means of three experiments (25, 50, or 100 width measurements for total body, BWM, and muscle cells, respectively, were made from n = 25 animals per strain in every experiment). Error bars indicate the standandard error of the mean. All differences between values at a given time point were statistically significant (p < 0.001 for A; p < 0.05 for B and C), except the BWM widths between *fhod-1(Δ)* and *cyk-1(ts)* at 0 hrs, which were not significant (p > 0.05).

**Figure S2.**
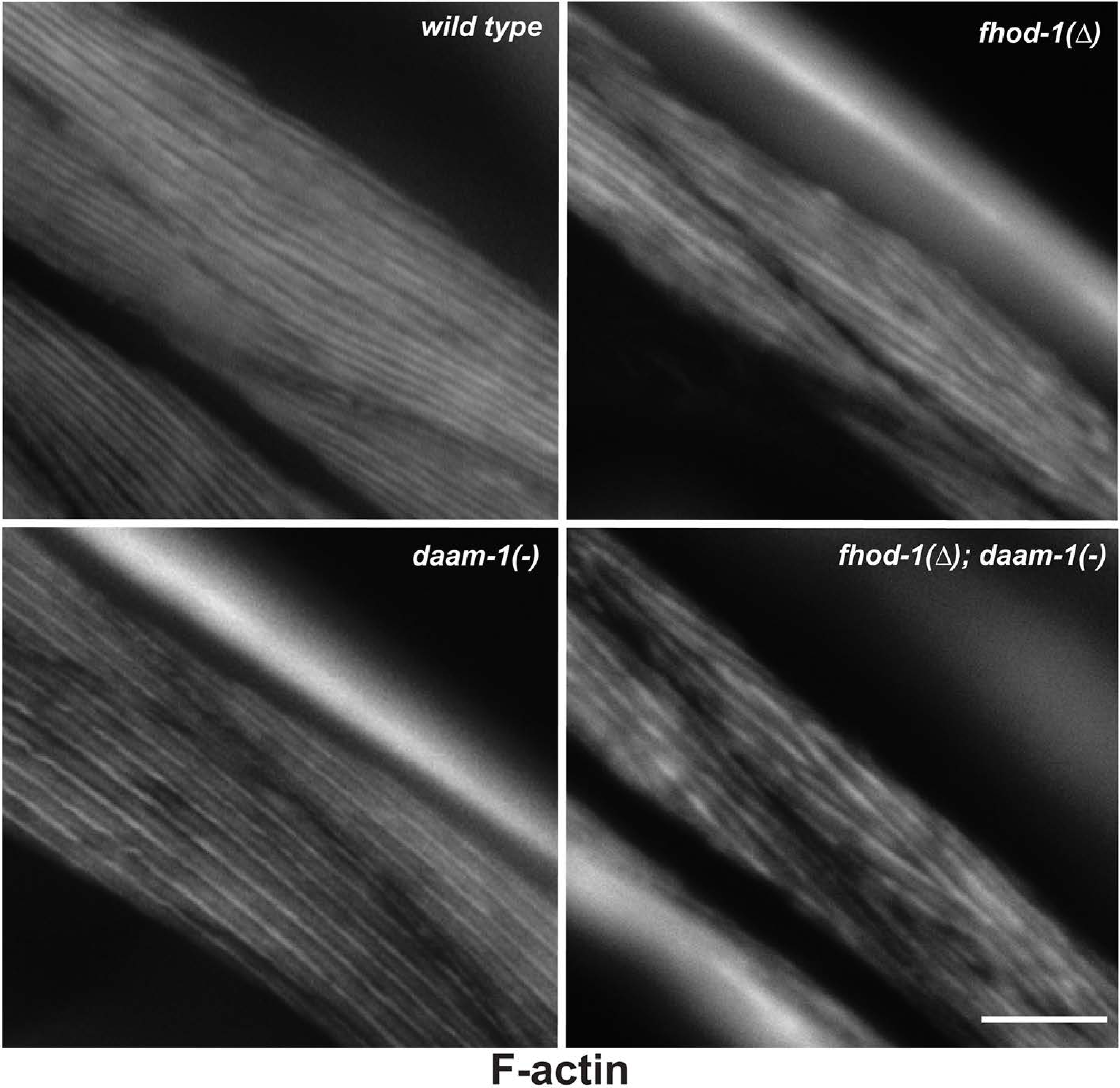
Absence of DAAM-1 did not cause any gross BWM defects. Dorsal views of adult worms of the indicated genotypes stained with fluorescently-labeled phalloidin to observe F-actin. Scale bar, 20 µm. There were no gross BWM defects observed due to absence of DAAM-1 compared to wild-type, or with a combined absence of both DAAM-1 and FHOD-1 compared to absence of FHOD-1, alone.

**Figure S3.**
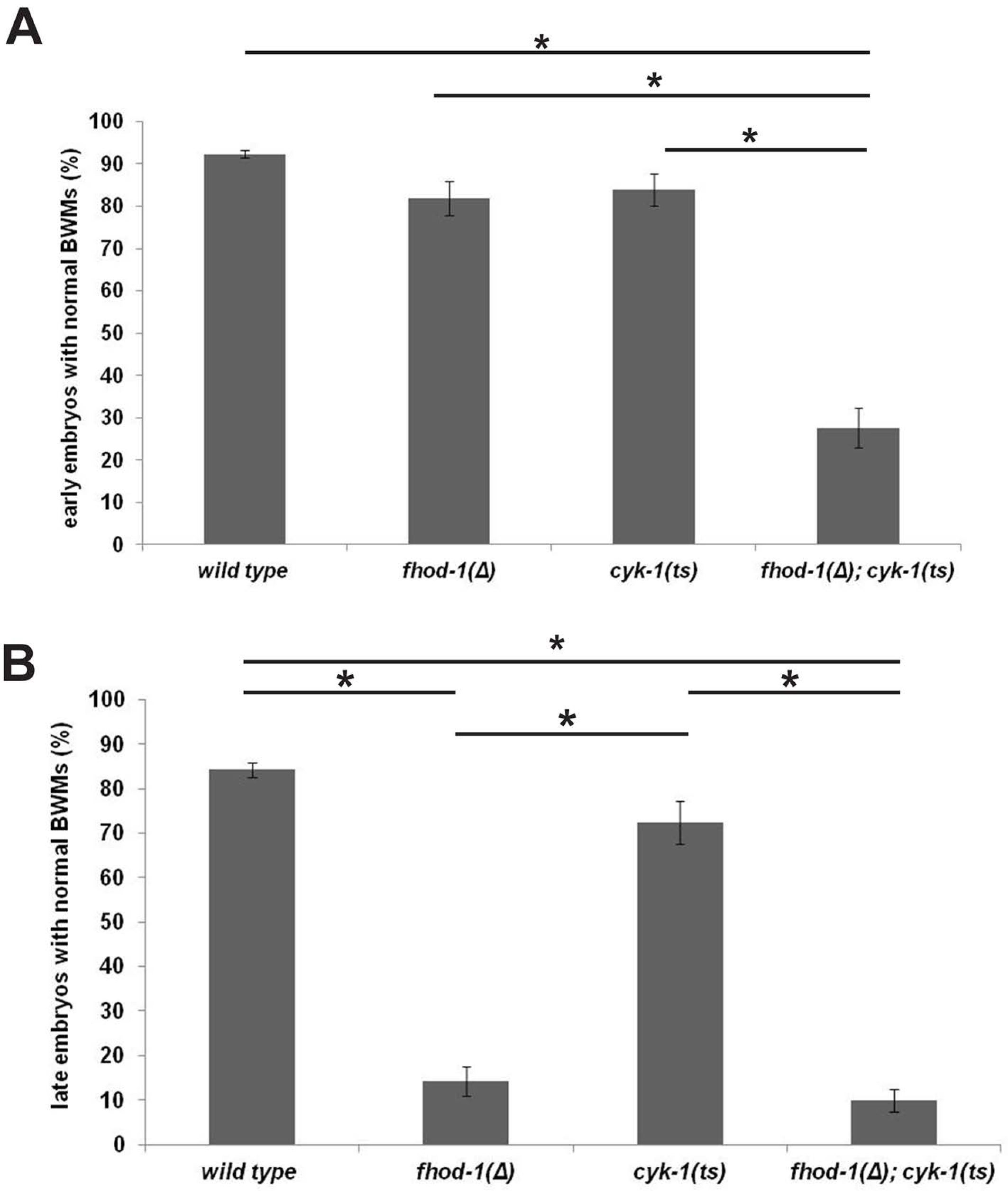
Embryonic *cyk-1(ts)* mutants at permissive temperature. Embryos expressing GFP-myosin were maintained at permissive temperature before staining with fluorescently-labeled phalloidin to observe F-actin. Graphs show percentage of embryos with normal BWMs for early embryos (A) and late embryos (B), based on the phenotypic categorization shown in Fig. 4. Graphs depict the average of the means of three experiments (n = 35 or 70 embryos for early and late embryos, respectively, per strain in every experiment). Error bars indicate the standandard error of the mean. (*) indicates p < 0.01. All other differences in pairwise comparisions were statistically insignificant (p > 0.05).

**Figure S4.**
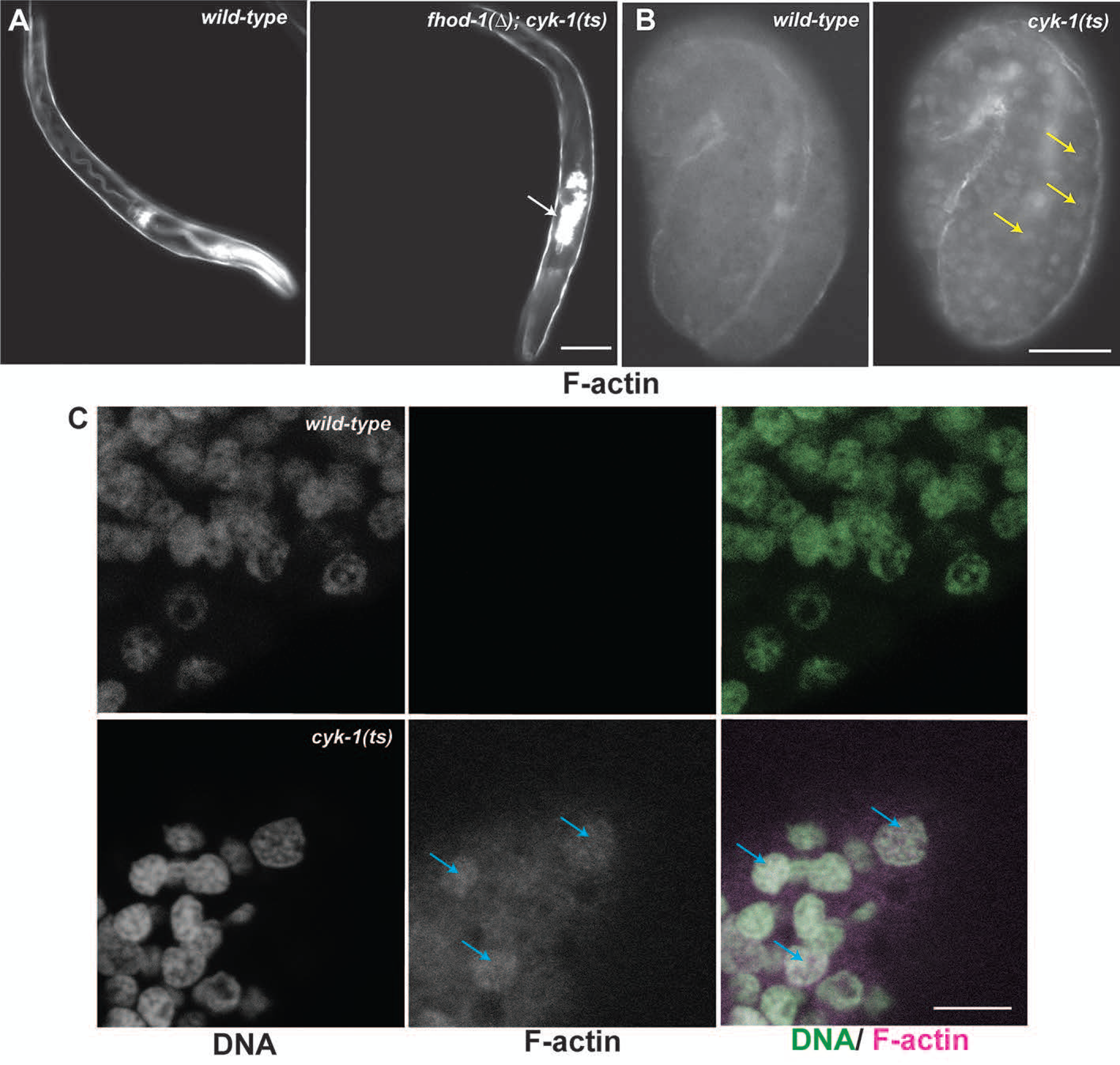
Detached pharynx and nuclear actin phenotypes observed in *cyk-1(ts)* mutants. (A) L1 larvae of wild-type and *fhod-1(Δ); cyk-1(ts)* double mutant stained with fluorescently-labeled phalloidin showing normal pharynx and a pharynx that has detached from the mouth (white arrow), respectively. Detached pharynx is a phenotype previously observed with simultaneous loss of CYK-1 and FHOD-1 due to deletion mutations or RNAi (Mi-Mi et al., 2012). Scale bar, 20 µm. (B) Early stage embryos of wild-type and *cyk-1(ts)* grown under restrictive conditions before being stained with fluorescently-labeled phalloidin to observe F-actin show *cyk-1(ts)* embryos have F-actin-rich round bodies (yellow arrows), while no such staining is observed in wild type embryos. Scale bar, 20 µm. (C) Confocal images show round F-actin bodies in *cyk-1(ts)* embryos are nuclei, based on their counter-staining with DAPI (arrows). Scale bar, 4 µm.

**Figure S5.**
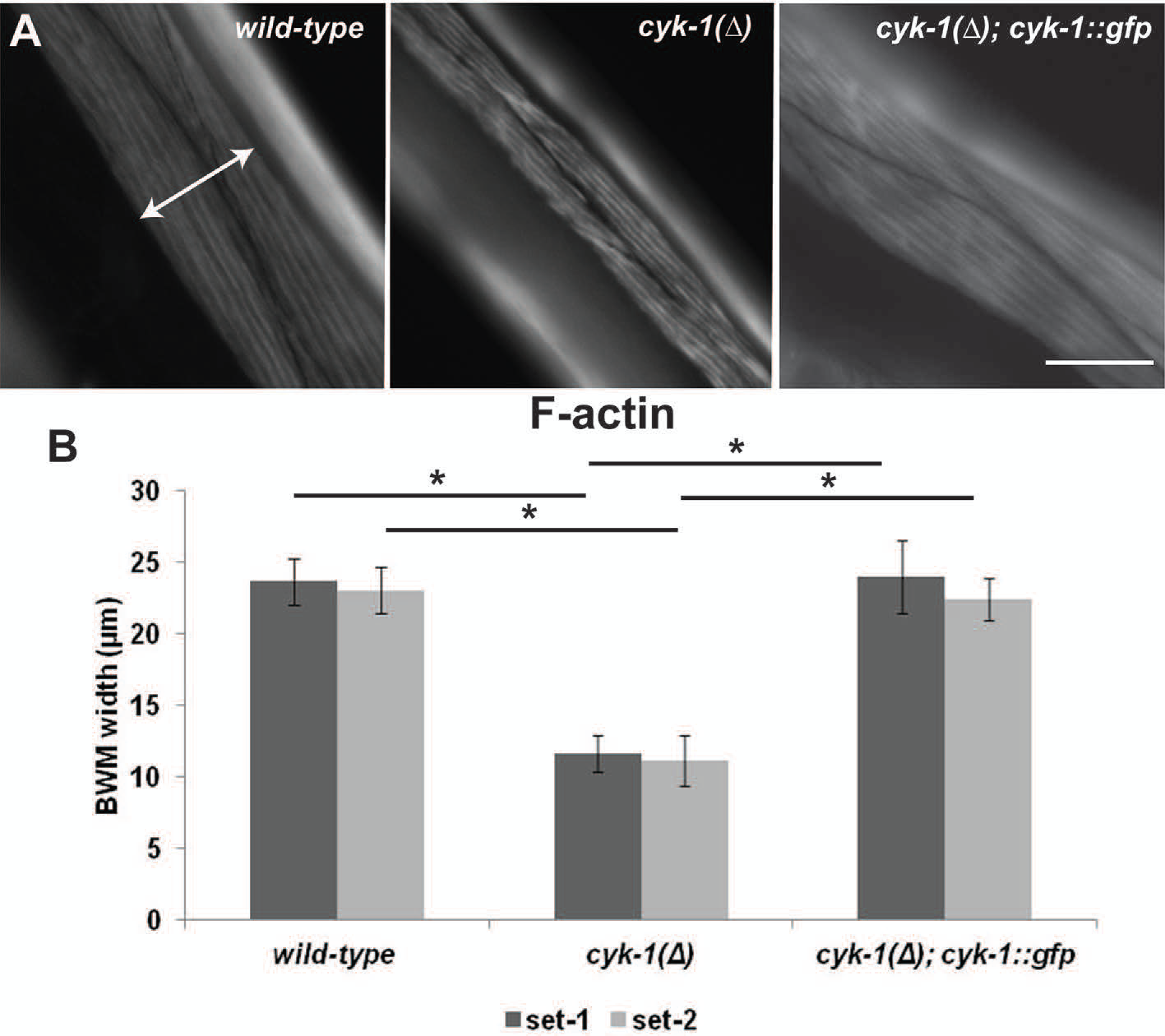
*cyk-1::gfp* rescues BWM width of *cyk-1(Δ)* mutants. (A) Dorsal views of adult worms of the indicated genotypes stained with fluorescently-labeled phalloidin to observe F-actin. Scale bar, 20 µm. (B) Meaurement of BWM widths (as shown in A, white double arrows) demonstrates normal BWM size is restored in *cyk-1(*Δ) animals with a *cyk-1::gfp* transgene integrated into the genome at an ectopic site. Shown are the means for two independent experiments (n = 11 animals for each genotype for each experiment, with 2 BWM width measurements per animal). Error bars indicate one standandard deviation. (*) indicates p < 0.01. Differences in all other pairwise comparisons were not statistically significant (p > 0.05). *cyk-1::gfp* is able to rescue the BWM width in *cyk-1(Δ)* mutants.

**Figure S6.**
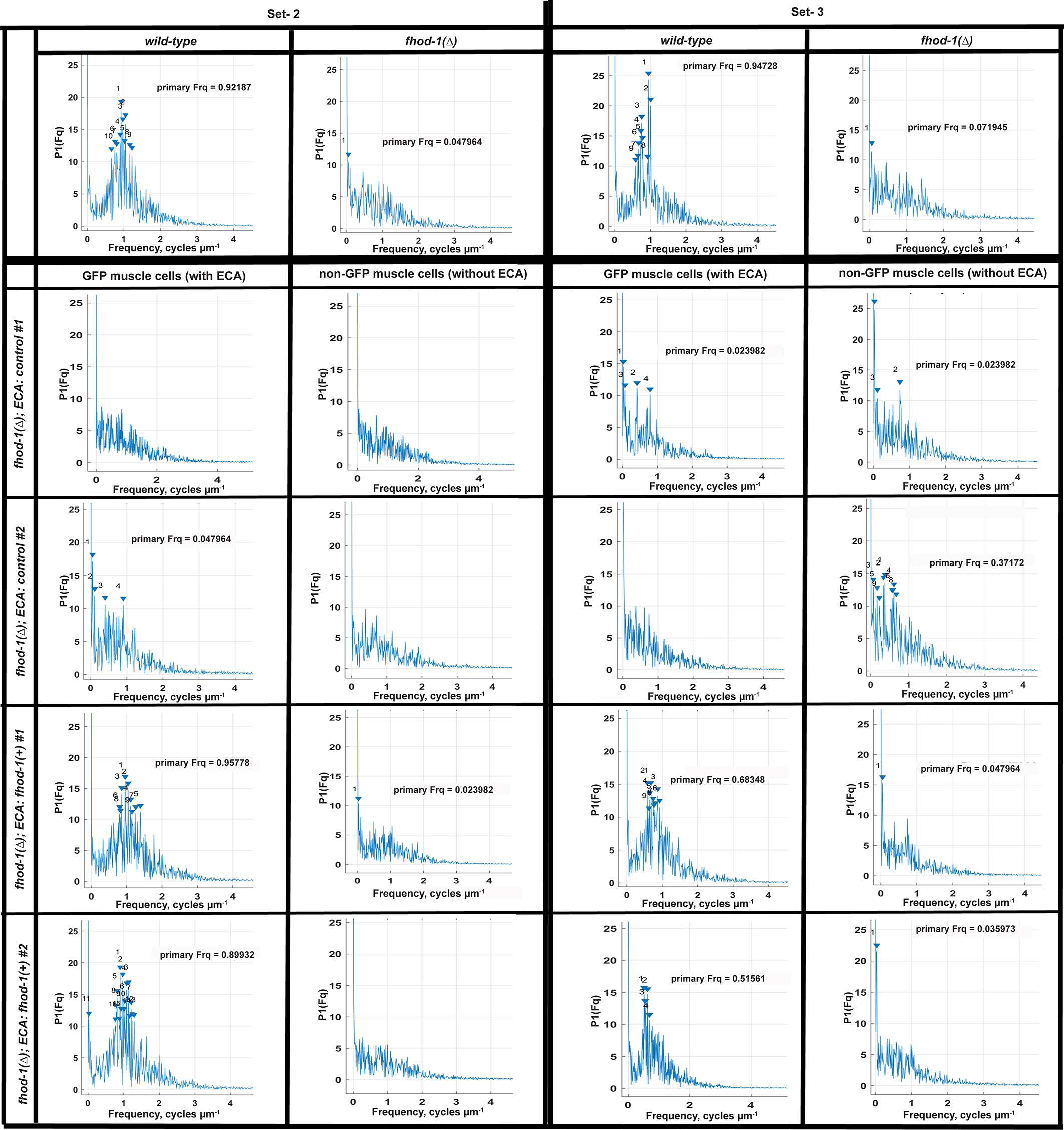
FHOD-1 plays a cell autonomous role in rescuing DB morphology. DB spacing was quantitatively analyzed for two independent sets of animals by performing FFT on concatenated intensity profiles of ATN-1 immunostain along all striations within single BWM cells (one ECA-containing BWM cell and one non-ECA-containing BWM cell per animal for transgenic lines, and one BWM cell for control strains; n = 10 animals per strain), similar to as done for Fig. 6. Amplitude spectra for DBs in wild type BWM cells or BWM cells that inherited a *fhod-1(+)-*containing ECA show clustering of peaks near a frequency of 0.8-1.0 µm^−1^, whereas amplitude spectra peaks did not cluster near any particular frequency for *fhod-1(*Δ) BWM cells that inherited no ECA, or inherited an ECA lacking *fhod-1(+)*.

**Figure S7.**
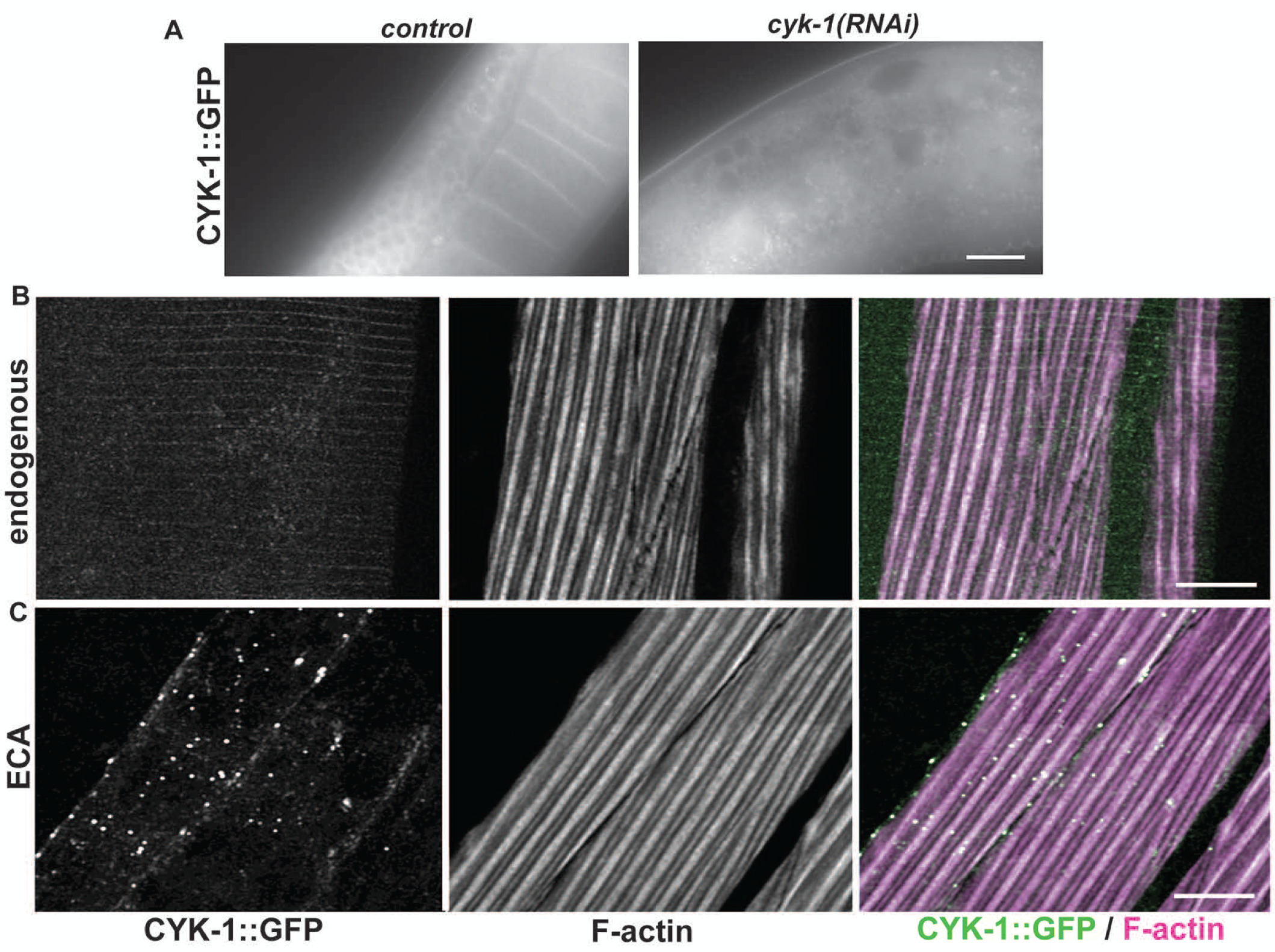
GFP-tagged CYK-1 localizes to the gonad but does not localize to DBs in BWM. (A) Worms where endogenous *cyk-1* was tagged with GFP were treated with control and *cyk-1(RNAi)* for 5 days. CYK-1::GFP expressed in the germline in control animals and this is absent in *cyk-1(RNAi)* animals. (B, C) Maximum intensity projections (MIP) of dorsal views of adult worms expressing CYK-1::GFP from (B) the endogenous *cyk-1* gene tagged with GFP or (C) from an ECA, and stained with fluorescently-labeled phalloidin. (B) Worms where the endogenous *cyk-1* gene is tagged with GFP show no CYK-1::GFP localization to DBs or any enrichment at all in BWM. (C) Worms with the ECA show punctuate CYK-1::GFP in F-actin-rich BWM, possibly due to over-expression, but show no localization of CYK-1::GFP to a pattern resembling DBs. Scale bars, 10 µm.

